# An aberrant protamine ratio is associated with decreased H4ac levels in murine and human sperm

**DOI:** 10.1101/2024.08.11.606797

**Authors:** Alexander Kruse, Simon Schneider, Gina Esther Merges, Andreas Christian Fröbius, Ignasi Forné, Axel Imhof, Hubert Schorle, Klaus Steger

## Abstract

Protamine 2 (*Prm2/PRM2*), together with Protamine 1 (*Prm1/PRM1*), constitute the two protamines found in both murine and human sperm. During spermiogenesis in haploid male germ cells, chromatin undergoes significant condensation, a phase in which most histones are replaced by a species-specific ratio of these two protamines. Altered PRM1/PRM2 ratios are associated with subfertility and infertility in both male mice and men. Notably, during histone-to-protamine exchange a small fraction of histones remains (ranging from 1% to 15%) bound to DNA. The regulatory roles of these residual histones, governed by post-translational modifications (PTMs), play a pivotal role in spermatogenesis, particularly in chromatin remodeling and epigenetic regulation of genes during sperm differentiation or even in early embryogenesis.

In this study, utilizing a *Prm2*-deficient mouse model and conducting an analysis of sperm samples from men exhibiting either normozoospermia or atypical spermiograms, we observed alterations in the methylation and acetylation profiles of histones H3 and H4. Subsequent in-depth analysis revealed that discrepancies in protamine ratios do not significantly influence the post-translational modifications (PTMs) of histones in testicular sperm. In epididymal sperm these altered protamine ratios are associated with a reduction in the acetylation levels of histone H4 (H4ac), a phenomenon consistent across both murine and human samples. In particular, H4K5ac and H4K12ac were identified as the two modifications that appear to decrease as a result of reduced *Prm2*/*PRM2* levels. Our findings reveal that Protamine 2 is necessary for the maintenance of specific histone PTMs, such as acetylation, which is essential for proper spermatogenesis and particularly for chromatin remodeling.

## Introduction

Spermatogenesis, the intricate process of male germ cell formation, involves the transformation of diploid spermatogonia into mature haploid spermatozoa through stages of mitosis, meiosis and spermiogenesis (Bergmann, 2006; Hess and de Franca, 2008). Spermiogenesis is marked by extensive chromatin restructuring, characterised by the sequential replacement of histones by protamines. This results in hypercondensation of the paternal chromatin, safeguarding DNA integrity and facilitating transcriptional silencing (Steger, 1999; Rathke *et al*., 2014; Bolcun-Filas and Handel, 2018).

Contrary to the majority of mammals, where sperm chromatin is packaged by a single protamine, primates, most rodent species, and a subset of other placental mammals uniquely possess two distinct protamines: protamine 1 (PRM1) and protamine 2 (PRM2) (Ammer *et al*., 1986; Oliva, 2006; Balhorn, 2007). In both mice and humans, the genes of protamine 1 (*Prm1/PRM1*) and protamine 2 (*Prm2/PRM2*) are located alongside the gene for transition protein 2 (*Tnp2/TNP2*) and the sequence of *gene 4* or "*Prm3*" within a conserved gene cluster on chromosome 16 (Reeves *et al*., 1989; Oliva, 2006). Diverging from protamine 1, protamine 2 is expressed as a precursor (pre-PRM2), which, upon binding to DNA, is sequentially cleaved at its N-terminal domain, resulting in the mature protamine 2 (mPRM2, henceforth referred to as PRM2) (Yelick *et al*., 1987; Oliva and Dixon, 1991). Protamines are characterised by their small size, a high number of arginine residues and the associated strong positive charge in contrast to the bound negatively charged DNA, and the ability to form disulphide bridges, mediated by cysteine-rich domains (Balhorn, 1982; Balhorn, 2007; Jodar and Oliva, 2014). Comparison of PRM1/PRM2 expression revealed species-specific ratios of 1:1 in men and 1:2 in mice (Corzett *et al*., 2002; de Mateo *et al*., 2009). Several studies, including mouse models and research on men with impaired fertility, have demonstrated a link between deviations in the protamine ratio and male subfertility and infertility (Cho *et al*., 2003; Aoki *et al*., 2005; Torregrosa *et al*., 2006; Steger *et al*., 2008; Ni *et al*., 2016; Schneider *et al*., 2016; Merges *et al*., 2022).

In addition to protamines, a small fraction of histones, characterized by specific posttranslational modifications (PTMs) remains bound to the DNA following the histone-protamine exchange (Tanphaichitr *et al*., 1978; Brunner *et al*., 2013; Chioccarelli *et al*., 2020). Specific genomic regions such as gene regulatory regions, retain their histones including PTMs and can influence gene expression in the early embryo (Guerrero-Bosagna and Skinner, 2014; Chioccarelli *et al*., 2020). Various histone PTMs contribute to chromatin remodelling during spermiogenesis in mice and men. In particular, the hyperacetylation of H4, which leads to reduced DNA-histone interaction, is one of the key modifications of the lysine residues of histones in this process (Bao and Bedford, 2016; Ketchum *et al*., 2018). Besides acetylation, several other histone PTMs can be found in the course of spermatogenesis, which are shown in the study by Luense et al. (Luense *et al*., 2016). This study highlighted a significant conservation in the PTMs of histones H3 and H4 across murine and human sperm. Particularly noteworthy were methylation (me1-3) and acetylation (ac) of lysine residues. Given that the sperm samples analysed in this study were normozoospermic, it raised intriguing questions about how the PTMs of H3 and H4 might differ in samples that deviate from this norm, and whether such variations could correlate with changes in the protamine ratio.

Therefore, in our study, we conducted a comparative analysis of changes in methylation and acetylation of histones H3 and H4 in murine and human sperm samples characterized by impaired spermatogenesis, reduced fertility potential (subfertile/infertile), and altered protamine ratios relative to their normal (fertile) counterparts, utilizing proteomic and immunodetection techniques. The murine samples used in this study were derived from the *Prm2*-deficient mouse model (*Prm2* mice = *Prm2*^+/+^, *Prm2*^+/-^ and *Prm2*^-/-^) established and analyzed by Schneider *et al*. (Schneider *et al*., 2016). In this model, heterozygous males retained fertility, whereas *Prm2^-/-^* mice displayed infertility. For human samples, we utilized fertile spermatozoa from ejaculates with normozoospermia and subfertile spermatozoa from ejaculate samples demonstrating abnormal protamine ratios and atypical spermiograms (manifestations including oligo-, astheno-, teratozoospermia or combinations thereof).

We found that PTMs of histones in murine testicular sperm are unaffected by either aberrant protamine ratio (*Prm2*^+/-^) or *Prm2*-deficiency (*Prm2*^-/-^). For epididymal sperm of *Prm2* mice, not only different levels of H3 and H4 but also different global modification levels of certain PTMs, especially for H4ac (H4K5ac, H4K8ac, H4K12ac and H4K16ac), were identified between the genotypes. Decreased H4ac levels found in *Prm2*^+/-^ and *Prm2*^-/-^ were also detected in subfertile human spermatozoa with aberrant protamine ratio and could be attributed to lower levels H4K5ac and H4K12ac. The comparable results between *Prm2*^+/-^ mice and sperm from subfertile men suggest that significant deviations in protamine ratios can also have epigenetic effects in form of altered global modification levels of specific core histone PTMs.

## Materials and methods

### Ethical approval

#### Mouse model

Mouse samples were obtained from the B6.B6D2F2-Prm2^em1Hsc^ strain, registered in the Mouse Genome Informatics database MGI:5760133 (Blake *et al*., 2021). Animals were generated and characterized according to the methodologies outlined previously (Schneider *et al*., 2016). These mice were housed under specific pathogen-free (SPF) conditions at the animal facility of the Biomedical Research Centre Seltersberg (BFS). During the retention period of the individual animals of at least eight weeks, no experiments were carried out. Sample collection was performed post-mortem, in accordance with the approval granted under JLU registration number 719_M (genetic engineering file number: IV44-53r30.03.UGI118). All personnel involved in this study possessed valid animal-handling certifications.

#### Human samples

All human samples used in this study were approved for research by the patients, the Clinic and Polyclinic for Urology, Paediatric Urology and Andrology of the UKGM (Ethic approval, Ref. No.: 95/04 and 146/06) and the Hormone and Fertility Centre of the Ludwig Maximilian University of Munich (LMU) (Project number: 23-0684). Semen analyses were conducted by qualified personnel in accordance with the protocols outlined in the WHO Laboratory Manual for the Examination and Processing of Human Ejaculate (World Health Organization, 2010). Ejaculate samples displaying deviations from standard parameters were identified based on key spermiogram metrics, namely concentration, morphology and motility. Two distinct cohorts were established for the study: The first, designated as “fertile”, comprised ejaculate samples that conformed to normozoospermic standards, while the second group, labelled “subfertile”, included samples exhibiting deviations in one or more primary parameters (oligo, astheno-, teratozoospermia or their combinations).

### Antibodies

Primary Antibodies: Rabbit polyclonal anti-Histone H3 antibody (Abcam, Cambridge, UK, cat#ab1791, dilution: IHC/IF/WB 1:500/1:1000), rabbit polyclonal anti-Histone H4 antibody (Abcam, Cambridge, UK, cat#ab7311, dilution: IHC/IF/WB 1:500/1:1000), mouse monoclonal anti-Histone H3 trimethyl K27 (H3K27me3) antibody (Abcam, Cambridge, UK, cat#ab6002, dilution: IHC/IF/WB 1:200/1:500), rabbit polyclonal anti-Histone H3 dimethyl K36 (H3K36me2) antibody (Abcam, Cambridge, UK, cat#ab9049, dilution: IHC/IF/WB 1:200/1:500), rabbit polyclonal anti-Histone H3 trimethyl K4 (H3K4me3) antibody (Abcam, Cambridge, UK, cat#ab8580, dilution: IHC/IF/WB 1:200/1:500), mouse monoclonal anti-Histone H3 dimethyl K9 (H3K9me2) antibody (Abcam, Cambridge, UK, cat#ab1220, dilution: IHC/IF/WB 1:200/1:500), rabbit polyclonal anti-Histone H3 monomethyl K79 (H3K79me1) antibody (Abcam, Cambridge, UK, cat#ab2886, dilution: IHC/IF/WB 1:200/1:500), rabbit polyclonal anti-Histone H3 trimethyl K79 (H3K79me3) antibody (Abcam, Cambridge, UK, cat#ab2621, dilution: IHC/IF/WB 1:200/1:500), rabbit monoclonal anti-Histone H4 acetyl K5 (H4K5ac) antibody (Abcam, Cambridge, UK, cat#ab51997, dilution: IHC/IF/WB 1:200/1:500), rabbit polyclonal anti-Histone H4 acetyl K8 (H4K8ac) antibody (Abcam, Cambridge, UK, cat#ab15823, dilution: IHC/IF/WB 1:200/1:500), rabbit polyclonal anti-Histone H4 acetyl K12 (H4K12ac) antibody (Abcam, Cambridge, UK, cat#ab46983, dilution: IHC/IF/WB 1:200/1:500) and rabbit monoclonal anti-Histone H4 acetyl K16 (H4K16ac) antibody (Abcam, Cambridge, UK, cat#ab109463, dilution: IHC/IF/WB 1:200/1:500), rabbit polyclonal anti-Histone H4 dimethyl K20 (H4K20me2) antibody (Abcam, Cambridge, UK, cat#ab9052, dilution: IHC/IF/WB 1:200/1:500), rabbit polyclonal anti-Histone H4 trimethyl K20 (H4K20me3) antibody (Abcam, Cambridge, UK, cat#ab9053, dilution: IHC/IF/WB 1:200/1:500).

Secondary Antibodies: Goat anti-mouse Ig/biotin (Dako, Hamburg, Germany, cat#E0433, dilution: 1:200), Goat anti-mouse IgG, Alexa Fluor 488 (Invitrogen, Darmstadt, Germany, cat#A11001, dilution: 1:10.000), Goat anti-rabbit Ig/biotin (Dako, Hamburg, Germany, cat#E0466, dilution: 1:200), Goat anti-mouse IgG, Alexa Fluor 568 (Invitrogen, Darmstadt, Germany, cat#A11004, dilution: 1:10.000), Goat anti-rabbit IgG, Alexa Fluor 488 (Invitrogen, Darmstadt, Germany, cat#A11008, dilution: 1:10.000), Goat anti-rabbit IgG, Alexa Fluor 568 (Invitrogen, Darmstadt, Germany, cat#A11011, dilution: 1:10.000)

### Immunohistochemistry

After excising the testicular tissue, the tissue was fixed in Bouin’s solution at 4°C overnight, followed by a wash in 70% ethanol. Each testis was then bisected and embedded in paraffin. For immunohistochemistry (IHC) 5 µm thick sections were placed on glass slides, deparaffinized and rehydrated. The sections were treated with a solution of 25 mM DTT, 0.2% (v/v) Triton x-100 and 200 i.U./ml heparin in phosphate-buffered saline (PBS) for 1 minute at 37°C to increase antigen accessibility (van der Heijden *et al*., 2006). Subsequently, the sections were washed in 0.02 M PBS (pH 7.4), boiled for 20 min in sodium citrate buffer, treated with 3% H2O2 in methanol for 20 minutes, and blocked with 5% bovine serum albumin (BSA) in PBS for 20 minutes. Primary antibody incubations were carried out overnight at 4°C. This was followed by a one-hour incubation at room temperature with a biotinylated goat anti-mouse or goat anti-rabbit secondary antibody (Dako, Hamburg, Germany), using the Vectastain ABC Kit (Vector Laboratories, Burlingame, CA, USA). As chromogen for the immunoreaction, AEC (Dako, Hamburg, Germany) was used, which remained on the section at RT until the desired staining intensity was achieved. Finally, the sections were counterstained with haematoxylin and sealed with Faramount aqueous mounting medium (Dako, Hamburg, Germany).

### Collection of murine epididymal sperm

Epididymal sperm samples were collected from sexually mature male mice aged between 2 and 5 months. The epididymides were carefully exposed, followed by a swim-out procedure in phosphate-buffered saline (PBS) at pH 7.4. This involved making several incisions in the cauda epididymis and gently pushing the sperm out. The sperm count was then determined using a Neubauer haemocytometer.

### Immunofluorescence

Murine and human sperm were washed with 2% Triton x-100 in PBS and smears were prepared on glass slides and air dried. These sperm smears were initially subjected to decondensation using decondensation Mix I (containing 50 mM Dithiothreitol (DTT) in 0.1 M Tris-HCl) for 15 minutes, followed by incubation with decondensation mix II (10 mM Lithium diiodosalicylate (LIS) + 5 mM DTT in 0.1 M Tris-HCl) for up to 2 h. Decondensation steps were performed at room temperature for human sperm and 37°C for murine sperm. After washing the smears three times with PBS, sperm were fixed in 4% paraformaldehyde (in PBS) for 0.5 to 1 hour. Blocking was performed using 5% bovine serum albumin (BSA) + 2% Triton x-100 in PBS, followed by overnight incubation with the primary antibody at 4°C and a 2-hour incubation with the fluorescently labelled secondary antibody at room temperature, along with Hoechst 33342 (Sigma-Aldrich/Merck, Darmstadt, Germany, cat#14533, dilution: 1:1000) for nuclear staining. Negative controls were prepared without the primary antibody. Following mounting with Faramount aqueous mounting medium, representative images were captured using an Axioskop 2 mot plus with Axiocam MRc (Zeiss, Munich, Germany).

### Histone extraction from sperm

Acid extraction of histones from murine and human sperm was conducted based on a published protocol (Shechter *et al*., 2007). For human samples, 40 million sperm were used and for murine samples, sperm from both epididymides of a single animal were utilized. After washing the sperm with PBS, they were centrifuged (5 minutes at 4000 rpm, RT) to form a pellet. The pellet was resuspended in PBS supplemented with 50 mM DTT and maintained on a rotary mixer for 30 minutes at 4°C to facilitate lysis. The histones were solubilized using 0.2 M sulfuric acid. In this solution, the samples were subjected to ultrasound treatment using a Bioruptor Pico (Diagenode, Seraing, Belgium, cat#B01060010) for 10 cycles of 30 seconds "ON" and 30 seconds "OFF" at 4°C to shear the DNA. Subsequently, the samples were incubated overnight at 4°C on a rotator, followed by centrifugation for 30 minutes at 14,000 rpm at 4°C. The supernatant was transferred into protein low-binding tubes (Eppendorf, Wesseling-Berzdorf, Germany, cat#0030108116). Protein and histone precipitation was achieved with 100% trichloroacetic acid (TCA) for 2 hours at 4°C. After washing the protein/histone pellets several times with ice-cold 100% acetone, they were air-dried for 15 min, dissolved in 1x Laemmli buffer and the pH was adjusted with 2 M Tris. The samples were then briefly stored at -20°C for subsequent use in immunostaining, immunoblotting or mass spectrometric analysis.

### SDS-PAGE and Western blotting

Protein/Histone extracts were denatured by heating at 95°C for 5 minutes in 1x Laemmli buffer (Bio-Rad, Feldkirchen, Germany, cat#1610747) and subsequently separated via 15% sodium dodecyl sulphate-polyacrylamide gel electrophoresis (SDS-PAGE). The electrophoresis was conducted for approximately 1.5 hours at a constant voltage of 100 V until reaching the separation gel, followed by a continued run at a constant 140 V. The proteins were then transferred to 0.45 µm pore size polyvinylidene fluoride (PVDF) membranes (Millipore, Boston, MA, USA) using the wet transfer method with a Mini Trans-Blot Cell (Bio-Rad, Munich, Germany). Total protein content, which was used for normalisation, was determined using the REVERT 700 Total Protein Stain kit (LI-COR, Lincoln, NE, USA, cat#926-11016). Subsequently, non-specific binding sites were blocked by incubating the membrane for 1 hour at room temperature with Intercept (PBS) blocking buffer (LI-COR, Lincoln, NE, USA, cat#927-76003). Incubation with the primary antibody (diluted 1:500 in blocking solution) was carried out overnight at 4°C. The following day, after multiple washes with PBST (PBS + 0.1% Tween 20), the membranes were incubated with a fluorescently labelled secondary antibody (1:10,000 dilution) for 1 hour at room temperature. Finally, the immunofluorescent signals were documented and analysed using the LI-COR imaging system Odyssey Fc 2800. The signal bands were quantified using the instrument’s software and normalized against total protein stain signals (Image Studio 5.2, LI-COR).

### Histone PTM Analysis via mass spectrometry

Histones were separated using a 16% polyacrylamide gel, followed by processing with propionic anhydride and trypsin, as previously described with slight modifications (Völker-Albert *et al*., 2018b). For liquid chromatography-mass spectrometry (LC-MS/MS) analysis, desalted peptides were injected in an Ultimate 3000 RSLCnano system (Thermo Fisher Scientific). These peptides were separated on a 15 cm analytical column (75 μm ID home-packed with ReproSil-Pur C18-AQ 2.4 (Dr. Maisch, Ammerbruch-Entringen, Germany) using a 50-min gradient ranging from 5 to 60% acetonitrile in 0.1% formic acid. The effluent from the HPLC was directly electrosprayed into a Q Exactive HF (Thermo Fisher Scientific) operating in the Top10 duty cycle. A survey full scan MS spectrum (from m/z 250–1400) was acquired with a resolution of R=60,000 at m/z 400 (AGC target of 3x10^6^). Ions with charges between+2 and +5 were sequentially isolated to a target value of 2x10^5^ (isolation window 2.0 m/z) and fragmented at 27% normalized collision energy. Typical mass spectrometric conditions included a spray voltage of 1.5 kV, no sheath and auxiliary gas flow; a heated capillary temperature of 250°C and an ion selection threshold of 33.000 counts. The raw data was analysed using Skyline (MacCoss Lab, (MacLean *et al*., 2010; Egertson *et al*., 2015)) as described previously (Völker-Albert *et al*., 2018a). Peak areas corresponding to the peptides and modifications of interest were exported to Microsoft Excel (CSV), and statistical analysis was conducted using GraphPad Prism version 8.4.3 for Windows (GraphPad Software, San Diego, CA, USA).

### RNA isolation and quantitative RT-PCR

RNA was extracted from testis tissue and sperm using the RNeasy Mini Kit (Qiagen, Hilden, Germany, cat# 74104), with tissue homogenization performed using the TissueLyser LT (Qiagen, Hilden, Germany) and an on-column DNase digestion conducted according to the manufacturer’s instructions. The concentration and purity of the isolated RNA were assessed using a NanoDrop spectrophotometer (Peqlab, Erlangen, Germany). First-strand cDNA synthesis was performed using the RevertAid H Minus First Strand cDNA Synthesis Kit (Thermo Scientific, Vilnius, Lithuania, cat# K1632) according to the manufacturer’s protocol, with the reaction carried out on a Thermal Cycler T100 (Bio-Rad). Quantitative real-time PCR (qPCR) analyses were performed on the CFX96 Touch system (CFX 96 Real-Time system + C1000 Touch thermal cycler, Bio-Rad), utilizing Bio-Rad CFX Manager 3.1 software and iQ SYBR Green Supermix (Bio-Rad, Feldkirchen, Germany, cat# 1708880). Each primer pair was validated using the NCBI primer blast tool (https://www.ncbi.nlm.nih.gov/tools/primer-blast/) and purchased from Thermo Fisher Scientific/Invitrogen. The sequences of the primers used for qPCR are shown in Supplementary Table SI. The cycling conditions were as follows: An initial denaturation step at 95°C for 5 minutes, followed by 40 cycles of 30 seconds at 95°C, 30 seconds at 60°C and 30 seconds at 72°C. The program concluded with a final extension at 72°C for 5 minutes and a melting curve analysis ranging from 55–95°C with a 0.5°C increment every 5 s. Relative mRNA expression levels were evaluated using the 2^-ΔΔ^Ct method and normalized to the expression of *ß-Actin* or *Catsper1* (reference gene) of the respective species. All experiments were performed in triplicate to ensure reproducibility and accuracy.

### Statistical analyses

Statistical analyses were conducted utilizing GraphPad Prism 8.4.3 for Windows (GraphPad Software). All quantitative results are presented as mean ± standard error of the mean (SEM). Appropriate statistical tests were employed to compare the groups (one-way analysis of variance (ANOVA) and Tukey’s test or multiple t-tests and Holm-Sidak method), including *Prm2*^+/+^, *Prm2*^+/-^, and *Prm2*^-/-^ mice, as well as human samples from fertile and subfertile individuals. Statistical significance was determined based on *P-*values, with a threshold for significance set at *P* < 0.05. The levels of significance were denoted as follows: not significant for *P* > 0.05; * for *P* ≤ 0.05; ** for *P* ≤ 0.01; *** for *P* ≤ 0.001; and **** for *P* ≤ 0.0001).

## Results

### Decreased levels of *Prm2/PRM2* are associated with reduced fertility

In order to determine the protamine ratios in *Prm2* mice and men, the relative expression of *Prm1/PRM1* and *Prm2*/PRM2 was analysed by qPCR. For this purpose, murine testicular material and sperm from ejaculates of men were used (Fig. 1 A and B). No significant differences could be detected between the genotypes of the *Prm2* mice for *Prm1*, whereas the relative expression of *Prm2* was halved in *Prm2^+/-^* compared to *Wt* and could not be detected in the infertile *Prm2^-/-^* mice (Fig. 1 A). Based on the values of the relative expression levels, the percentage of *Prm1* and *Prm2* in relation to the total expression of *Prm1* + *Prm2* could be determined and was as follows: *Wt*: *Prm1* = 43%, *Prm2* = 57%; *Prm2^+/-^*: *Prm1* = 70%, *Prm2* = 30%; *Prm2^-/-^*: *Prm1* = 100%, *Prm2* = 0%. The comparison of human fertile and subfertile spermatozoa showed a comparable reduction of *PRM2* as found in *Prm2^+/-^* mice and also no significant differences for *PRM1* (Fig. 1 B). The percentage distribution of protamines in men was as follows: Fertile: *PRM1* = 55%, *PRM2* = 45%; Subfertile: *PRM1* = 64%, *PRM2* = 36%.

**Figure 1.**
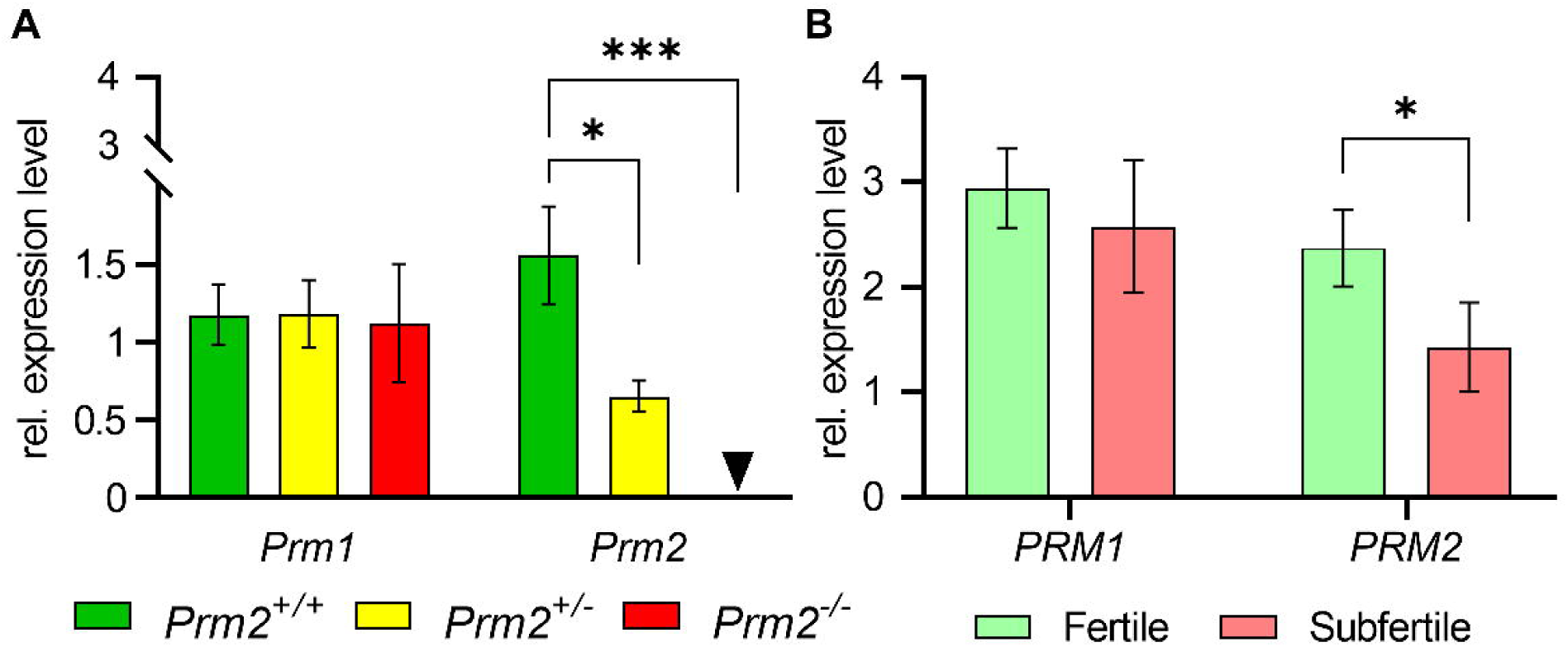
Differences in the relative expression of *Prm1/PRM1 and Prm2/PRM2*. **A**: Relative expression levels of *Prm1* and *Prm2* in testes of *Prm2* mice were determined using qPCR and normalized to *Catsper1* (ΔΔCt method). Mean values with corresponding standard errors (standard error of the mean [s.e.m.]) are shown for the different genotypes. Significant differences were determined using one-way analysis of variance (ANOVA) and Tukey’s test: *Prm2^+/+^/ Prm2^+/-^* * p = 0.0449; *Prm2^+/-^/ Prm2^-/-^* *** p = 0.0001; n = 10 per genotype. Arrowhead: The relative expression level for *Prm2* in *Prm2^-/-^*-testes was 0. **B**: Determination of the relative expression levels of *PRM1* and *PRM2* in human sperm was conducted using qPCR and normalized to β-actin (ΔΔCt method). Mean values with corresponding standard errors (s.e.m.) are shown for the fertile and subfertile group. The comparative statistical analysis was performed using multiple t-tests, with p-values for multiple testing adjusted using the Holm-Sidak method: *PRM2*: * p = 0.0407; Fertile n = 60; Subfertile n = 40.

It can be summarised that alterations in the protamine ratio are associated with reduced fertility in both mice and men, with the ratio changes observed in this study being specifically due to lowered *Prm2/PRM2* levels.

### Altered *Prm2* levels do not affect H3 and H4 expression patterns in testis but lead to changed histone amounts in epididymal sperm of *Prm2* mice

To assess the impact of *Prm2*-deficiency on the protein level of H3 and H4 during murine spermatogenesis, immunostaining was conducted (see Fig. 2 A-C and E-G). H3 exhibited nuclear staining in at least two distinct cell types across all twelve stages of spermatogenesis, with an absence of staining in elongated spermatids beyond step 12 (Fig. 2 A-D). For H4, regarding spermiogenesis, staining was only observed in elongating spermatids from steps 9-12. In addition, spermatogonia, as well as (pre-) leptotene and zygotene spermatocytes were positive for H4 (Fig. 2 E-H). Although the expression patterns of H3 and H4 differed, no genotype-specific variations could be discerned in testes.

**Figure 2.**
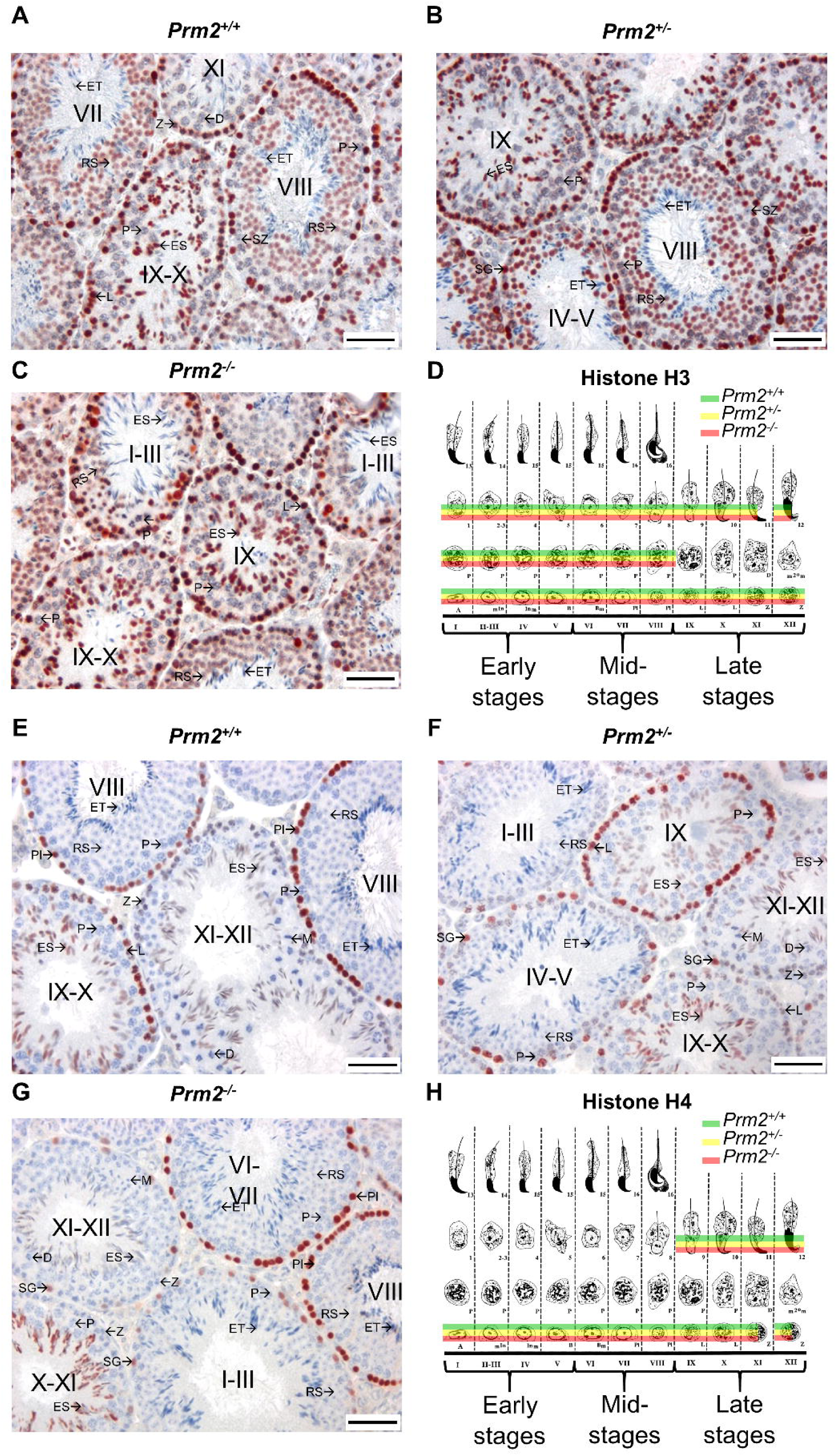
Histone H3 and H4 expression in murine testis of *Prm2*-deficient mice. Shown are representative images of the immunohistochemical detection of histone H3 (**A**-**C**) and H4 (**E**-**G**) in testis from *Prm2* mice and a schematic representation of the stage-specific expression patterns (**D** and **H**). A half overlay of cell types in the schematics indicates weak or partial staining (figure modified after (Russell *et al*., 1993; Endo *et al*., 2015)). Histone H3/H4 positive cells showed red staining and remaining nuclei were stained blue by counterstaining with haematoxylin. Stages of spermatogenesis are indicated by roman numerals. Stainings of five animals were analysed per genotype (n = 5); cell types: Spermatogonia type A (A); type A spermatogonia in mitosis (m^In^), intermediate spermatogonia in mitosis (In_m_), type B spermatogonia (B), type B spermatogonia in mitosis (B_m_) ◊ summarised as spermatogonia (SG), preleptotene spermatocytes (Pl), leptotene spermatocytes (L), zygotene spermatocytes (Z), pachytene spermatocytes (P), diplotene spermatocytes (D), secondary spermatocytes in meiosis (m2^°^m/M), 1-8 round spermatids (RS), 9-16 elongating (ES)/elongated (ET) spermatids; scale bar = 50 µm.

The effect of an altered protamine ratio on H3 and H4 expression in male germ cells after histone-protamine exchange was evaluated using immunostaining (Supplementary Fig. S1) and immunoblotting (Fig. 3 A and D). These investigations involved epididymal sperm from *Prm2* mice and human sperm from fertile and subfertile men. Immunofluorescent staining of histone H3 and H4 in epididymal sperm of *Prm2* mice was evident in nearly all sperm heads, with staining localized near the acrosome in the nuclear region of the sperm heads, covering approximately half of the nucleus (Supplementary Fig. S1 A and B). No significant differences were detected across genotypes. Deviating from the results of the publication on *Prm1* mice (Merges *et al*., 2022) the quantification using immunoblot analysis indicated that the detected signal intensities for both H3 and H4 varied significantly among the different genotypes of *Prm2* mice. In epididymal sperm of *Prm2* mice, H3 exhibited an increased signal for *Prm2^+/-^* compared to *Wt* and *Prm2^-/-^*, while for H4, sperm of *Prm2^-/-^* mice showed elevated signal values compared to *Wt* and *Prm2^+/-^* (Fig. 3 A-C). Similarly, to the murine staining, immunofluorescence analysis of histones H3 and H4 in human spermatozoa exhibited strong nuclear-associated staining, with no significant differences observed between fertile and subfertile samples (Supplementary Fig. S1 C and D). The results of the immunoblot analysis highlighted a significant disparity in histone quantities in human sperm, with H4 levels significantly exceeding those of H3 (Fig. 3 D-F). Intriguingly, there were no discernible differences in histone content between fertile and subfertile human sperm samples (Fig. 3 E and F).

**Figure 3.**
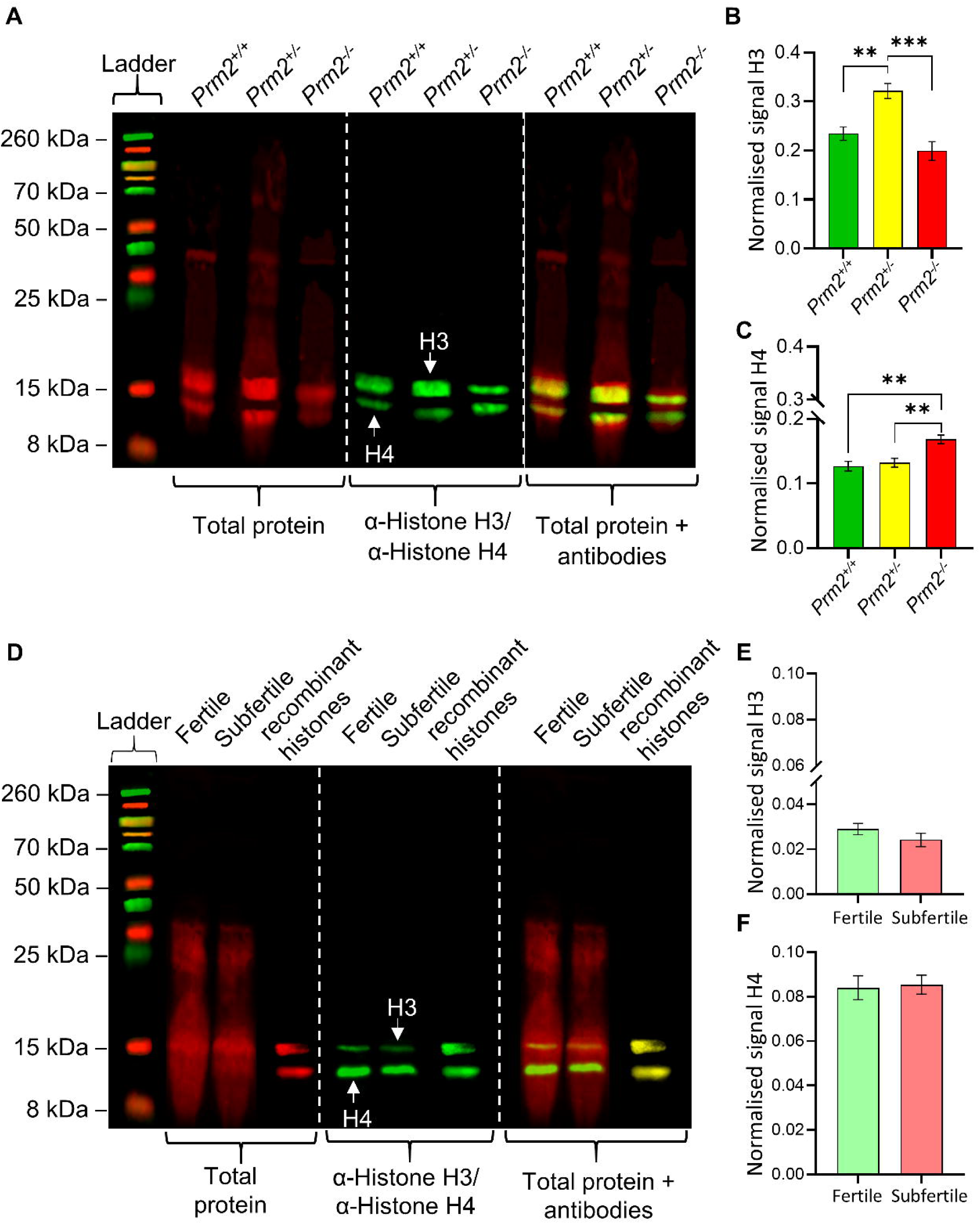
Varying H3 and H4 levels in murine and human spermatozoa. Histones H3 and H4 were analysed by immunoblot for epididymal sperm of *Prm2* mice (**A**-**C**) and spermatozoa from fertile and subfertile men (**D**-**F**). In addition to antibody-based detection, representative images also show the labelled total protein, which was used to normalise signal values of H3 and H4. Based on protein-ladder and recombinant histones, protein masses of H3 with ∼15 kDa and of H4 with ∼11 kDa could be assigned. Signal values of the bands for H3 (**B** and **E**) and H4 (**C** and **F**) were determined via Image Studio Ver. 5.2 software and normalised using the signal values of corresponding total protein. Shown are the mean values of the normalised signal with s.e.m. and significant differences were detected via single factor analysis of variance (ANOVA) and Tukey test. Histone H3: *Prm2^+/+^/ Prm2^+/-^* ** p = 0.0063; *Prm2^+/-^/ Prm2^-/-^* *** p = 0.0005; Histone H4: *Prm2^+/+^/ Prm2^-/-^* ** p = 0.0034; *Prm2^+/-^/ Prm2^-/-^* ** p = 0.0084; Per genotype n = 5.

These findings demonstrate that H3 and H4 are present in various cell types within murine testis and remain unaffected by changes in the protamine ratio. The presence of H3 and H4 in murine and human sperm is similar but not identical, with differing H3 and H4 levels observed in fertile sperm. Furthermore, it was demonstrated that loss of *Prm2* in mice can impact the overall H4 level in sperm. Sperm of heterozygous mice had an increased H3 level, whereas human subfertile sperm did not exhibit such changes in histone levels for H3 or H4, despite presumptions of an altered protamine ratio.

### PTM patterns of H3 and H4 differ between fertile and sub-/infertile sperm

Given the observed variations in the abundance of H3 and H4 between fertile and sub-/infertile sperm in mice, the subsequent phase of our study focused on comparing differences at the epigenetic level. Our goal was to evaluate the extent of modifications of H3 and H4 within murine and human sperm and to ascertain if reduced *Prm2* level might lead to changes in the frequency of specific post-translational modifications (PTMs). To achieve this, histone/protein extracts were initially subjected to mass spectrometric analysis. This technique facilitated the quantification of the percentage of various PTMs per peptide. Notably, across both histones (H3 and H4), the PTM percentages per peptide spanned a wide range, with some PTMs registering as low as 1% or less, while others exceeded 50% (Fig. 4).

**Figure 4.**
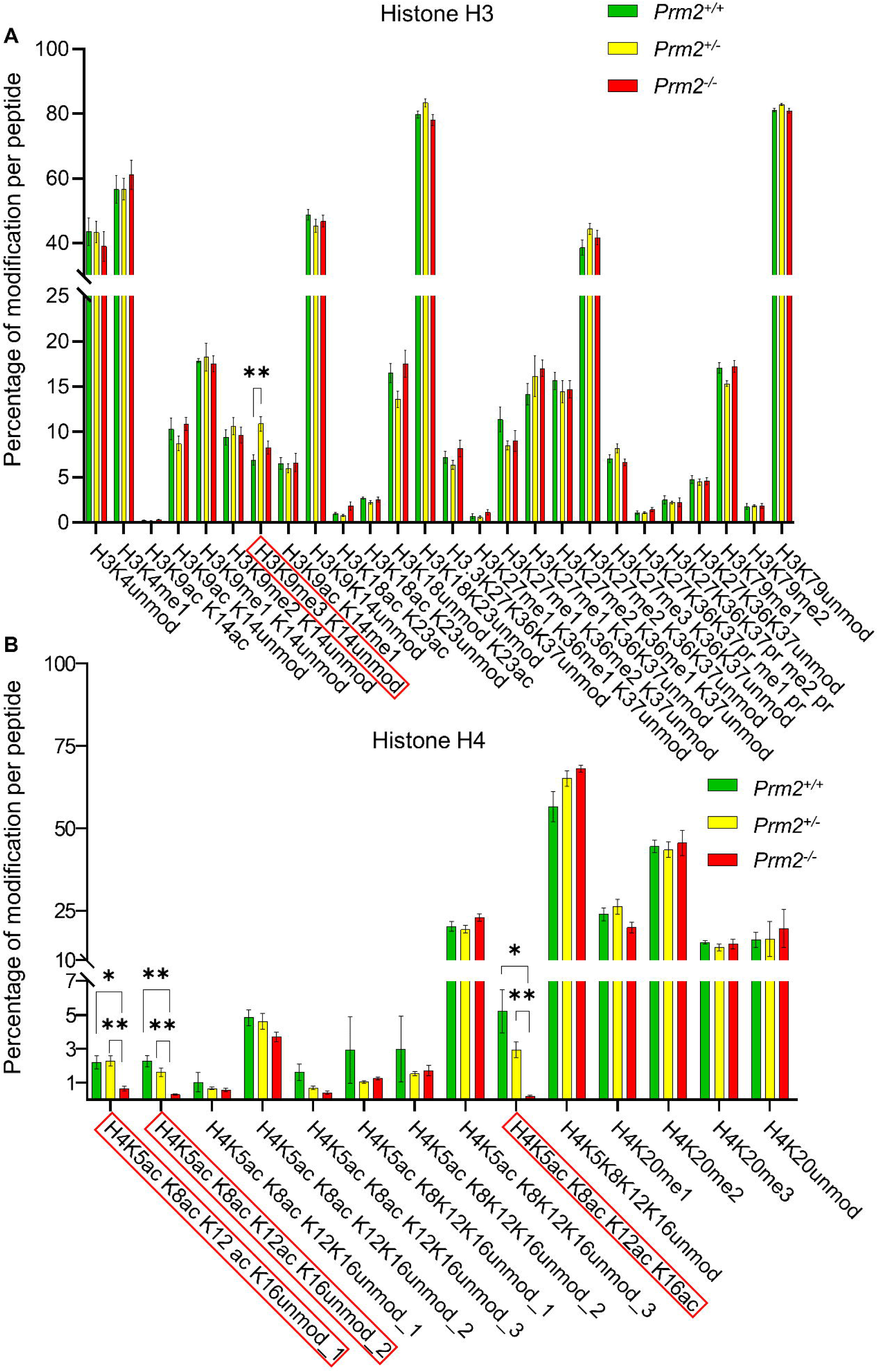
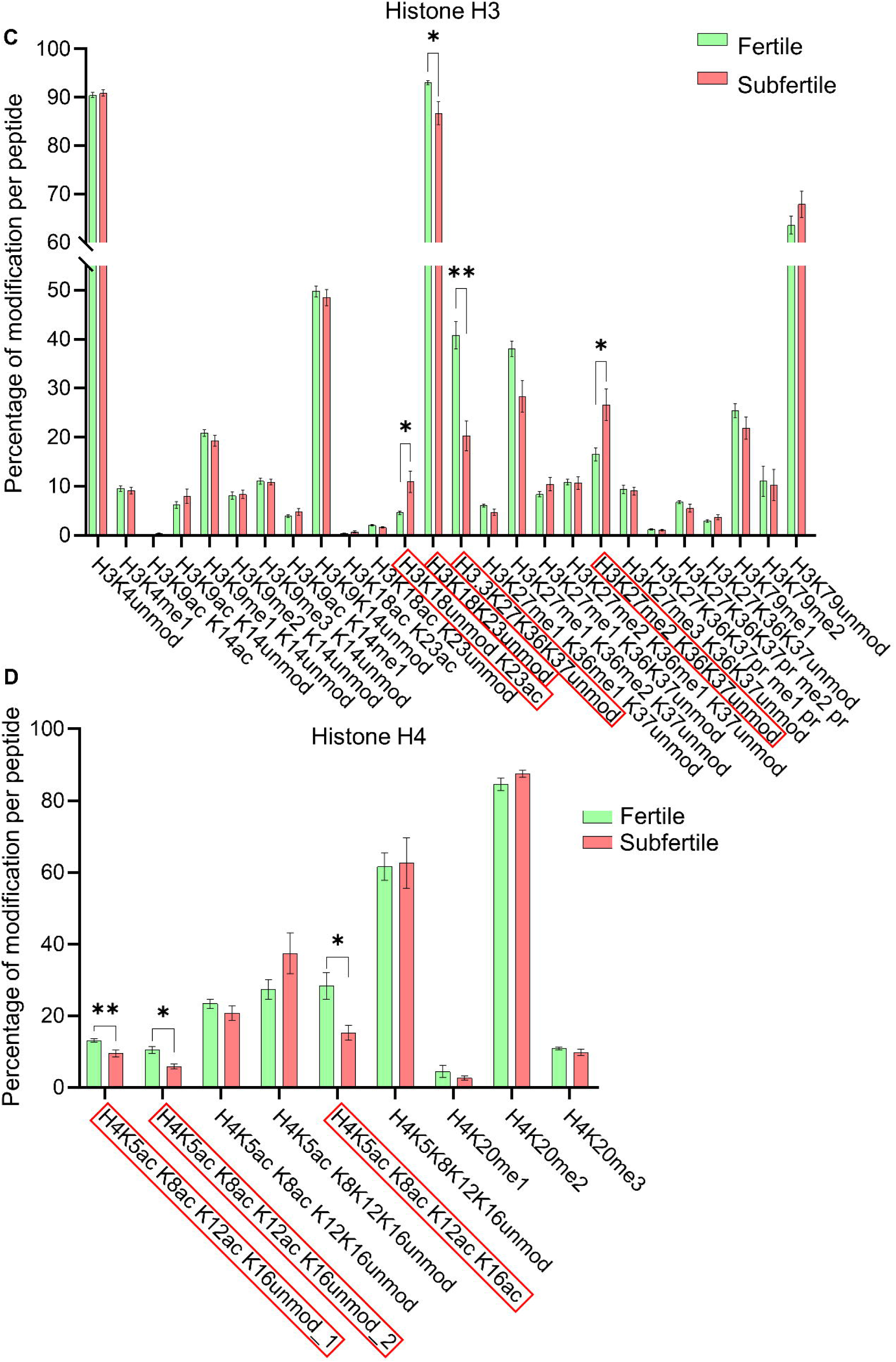
PTMs of H3 and H4 in sperm from *Prm2*^+/-^ mice and human subfertile sperm show similar changes. Using bottom-up LC-MS/MS, the histones extracted from epididymal sperm of *Prm2* mice and from human fertile/subfertile sperm were analysed for their PTMs (methylation and acetylation). Shown are the mean values of the percentage of the respective modifications per peptide with standard error (s.e.m.) for the core histones H3 (**A** and **C**) and H4 (**B** and **D**). Modifications shown are acetylation (ac) and methylation (me), which are listed after the respective lysine. If there are several unmodified (unmod) lysines, these are listed together. A numbering (e.g. _1) of the listed peptides including histone modifications represents one of the different peaks measured, which could be assigned to the same histone modifications, but differed minimally in intensity and retention time. The comparative statistical analysis was performed using multiple t-tests, whereby the p-values for multiple tests were adjusted using the Holm-Sidak method. In mice (**A** and **B**), epididymal sperm from eight animals (n = 8) per genotype group were analysed. The group size of human samples (**C** and **D**) was n = 20 for fertiles and n = 10 for subfertiles. **A**: Only H3K9me3 K14unmod showed a significant difference (red box) when comparing *Wt* and *Prm2^+/-^* ; H3K9me3 K14unmod: *Prm2^+/+^/Prm2^+/-^* : * p = 0.047; **B**: Histone H4 showed several significant differences (red boxes), especially with regard to acetylation; H4K5ac K8ac K12ac K16unmod_1: *Prm2^+/+^/Prm2^-/-^* : * p = 0.021; *Prm2^+/-^/Prm2^-/-^*: ** p = 0.0026; H4K5ac K8ac K12ac K16unmod _2: *Prm2^+/+^/Prm2^-/-^*: *** p = 0.0004; *Prm2^+/-^/Prm2^-/-^* ** p = 0.0023; H4K5ac K8ac K12ac K16ac: *Prm2^+/+^/Prm2^-/-^* : * p = 0.013; *Prm2^+/-^/Prm2^-/-^*: ** p = 0.00056. **C**: Significant differences (red boxes) between fertiles and subfertiles could be determined: H3K18unmod K23ac: * p = 0.011; H3K18K23unmod: * p = 0.031; H3.3K27K36K37unmod: ** p = 0.0027; H3K27me2 K36K37unmod: * p = 0.037; **D**: For H4, multiple acetylated peptides in particular showed significant differences (red boxes) when comparing the two groups (fertile vs. subfertile): H4K5ac K8ac K12ac K16unmod_1: ** p = 0.0093; H4K5ac K8ac K12ac K16unmod_2: * p = 0.02; H4K5ac K8ac K12ac K16ac: * p = 0.026.

For murine sperm, a single significant difference was identified in the genotype comparison for histone H3, in the trimethylation of H3K9 (H3K9me3 K14unmod) between *Wt* and *Prm2^+/-^* mice (Fig. 4 A, red box). Furthermore, no significant genotype-related variations were found for the histone variant H3.3 compared to the H3.1 variant (Fig. 4 A). In case of histone H4, significant genotype-based differences were observed in acetylation levels as highlighted in Figure 4 B (indicated by red boxes). These differences could be found in peptide H4 4-17, encompassing lysines K5, K8, K12 and K16, in terms of the proportion of modification per peptide. For H4K5ac K8ac K12ac K16ac, a gradational pattern was observed in the modification percentages corresponding to the decreasing *Prm2* level. Mean values were 5.21% for *Prm2^+/+^*, 2.94% for *Prm2^+/-^* and 0.22% for *Prm2^-/-^* (Fig. 4 B).

Unlike the murine results, an analysis of histone PTMs in sperm from fertile and subfertile men with altered protamine ratios revealed several significant differences in PTMs of histone H3 (Fig. 4 C). A significant increase in the modifications H3K23ac and H3K27me2 was observed in subfertile sperm (Fig. 4 C, red boxes). Additionally, a significantly reduced amount of the histone variant H3.3 (H3.3K27K36K37unmod) was detected in subfertile sperm samples (Fig. 4 C, red box). For histone H4, similarities with murine results were apparent, highlighting significant differences in PTM percentages for peptide H4 4-17 between fertile and subfertile group (Fig. 4 D, red boxes). These differences were characterised by reduced modification percentages in the subfertile group with altered protamine ratio, exemplified by H4K5ac K8ac K12ac K16ac with approximately 28% in fertile sperm and about 15% in subfertile sperm (Fig. 4 D, red box).

In summary, the PTMs of core histones H3 and H4 exhibit variations from the control group in both men and mice with altered protamine ratio (*Prm2*-deficiency), as shown by the mass spectrometric results.

### Altered Protamine ratios in sperm: Minor impact on H3 PTMs, similar H4 PTM abnormalities in human and murine samples

Guided by the findings from the mass spectrometric analysis, our subsequent research primarily focused on the following post-translational modifications (PTMs) of H3 and H4:

**H3K4me3, H3K9me2, H3K27me1, H3K27me3, H3K36me2, H3K79me1, H3K79me3 H4K5ac, H4K8ac, H4K12ac, H4K16ac, H4K20me2 and H4K20me3**

Initially, the status of each of these PTMs was examined in testicular tissue sections of *Prm2* mice (H3: Supplementary Fig. S2 and H4: Supplementary Fig. S3). Stainings showed no significant differences between genotypes in global modification levels of the listed PTMs in spermatids after step 8 of murine spermiogenesis (elongating and elongated spermatids).

Following preliminary investigations, similar analyses were not conducted on human testicular sections. This decision was due to the availability of only normal spermatogenesis (NSP) tissue samples and testis sections with spermatid arrests at the round spermatids (SDA) stage, which are unsuitable for assessing the influence of protamines due to the absence of cell types expressing protamines. Consequently, the focus for subsequent experiments shifted to the examination of epididymal sperm in *Prm2* mice and human sperm derived from ejaculate samples.

Through immunofluorescent staining of murine and human sperm targeting specific PTMs, as depicted in Figure 5, and supplemented by semi-quantitative Western blot analysis shown in Figure 6, we identified modifications that exhibited parallel changes in *Prm2^+/-^* or *Prm2*^-/-^ mice and subfertile human sperm in comparison to their respective control groups. Insignificant staining results can be found in the supplementary data (Supplementary Fig. S4 and S5). The only significant difference in H3 modifications was observed for H3K4 trimethylation (H3K4me3), which was decreased in subfertile human sperm as shown by Western blot analysis (Fig. 6 A, B, D and E). Corresponding immunofluorescent staining for H3K4me3 (Fig. 5) has shown that especially sperm with abnormal head morphology were positive for H3K4me3 but negative for the nuclear marker Hoechst, in both murine and human sperm. Further significant differences between the genotypes or between fertile and subfertile sperm could not be identified. In contrast, significant changes were observed in two selected PTMs of H4, regarding both sperm head staining and global modification levels. The acetylation level of H4K5 in *Prm2^-/-^* mice showed a significant decrease compared to the results of *Wt* and *Prm2^+/-^* mice, a trend also observed in the comparison of subfertile human sperm to the fertile group (Fig. 5, Fig. 6 A, C, D and F). H4K12ac emerged as another noteworthy PTM of H4. This modification exhibited a significant and progressive decrease from *Wt* to *Prm2^-/-^* sperm. Both immunofluorescent staining and Western blot analyses revealed weaker signals for *Prm2^+/-^* compared to *Wt* and nearly undetectable signals for *Prm2^-/-^* sperm (Fig. 5, Fig. 6 A and C). Subfertile human sperm displayed a significantly reduced level of H4K12ac compared to fertile sperm, as evidenced by immunofluorescence and Western blotting (Fig. 5, Fig. 6 D and F). Therefore, the results of the subfertile human sperm in particular were comparable with those of the *Prm2^+/-^* mice. However, the other investigated PTMs of H4 did not exhibit such distinct differences among the respective groups in either murine or human sperm (Fig. 5, Fig. 6).

**Figure 5.**
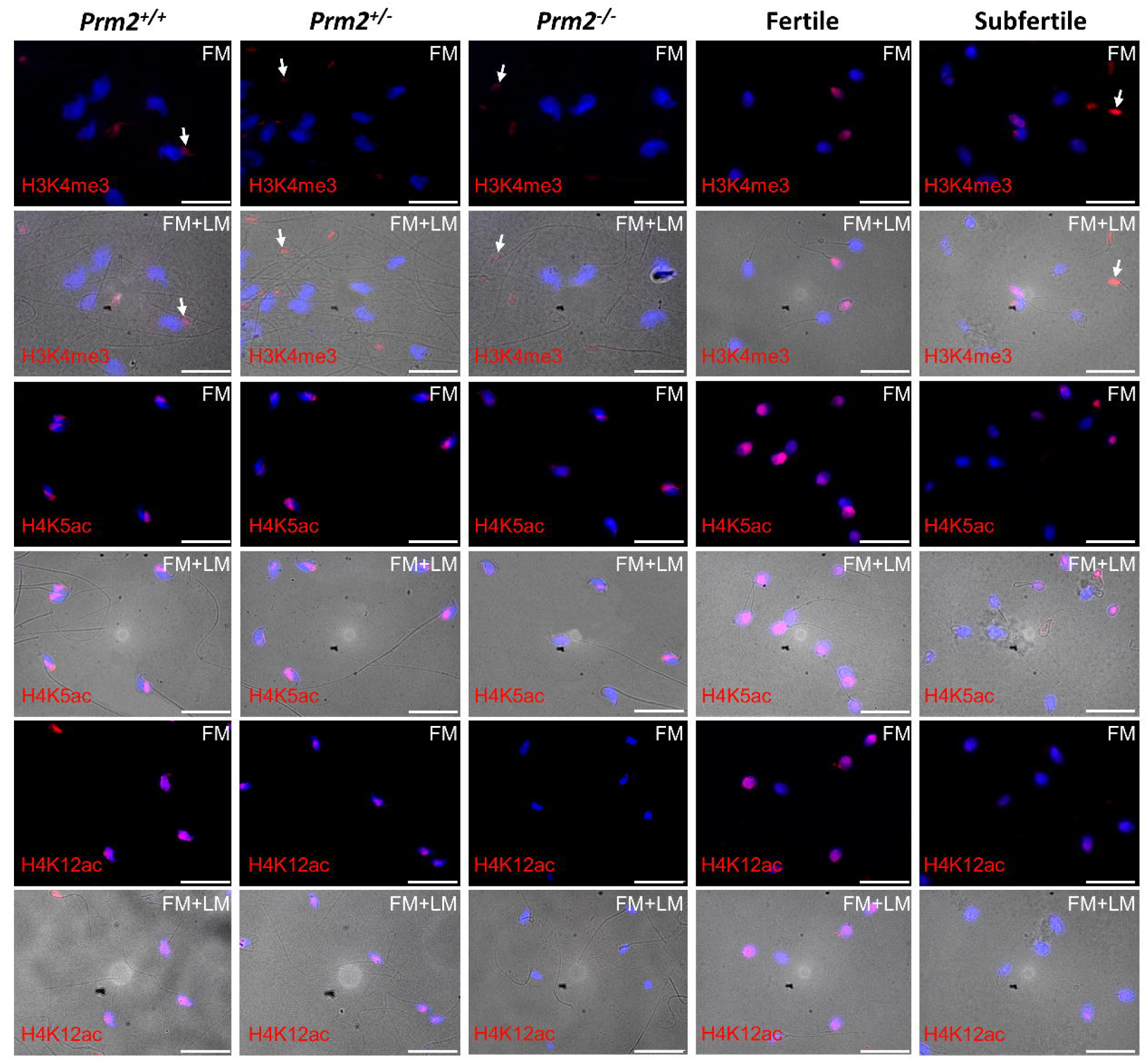
Decreasing acetylation of lysines K5 and K12 occurs in mice and men together with a deviating protamine ratio. For the detection of selected PTMs of H3 and H4, immunofluorescent stainings were performed on sperm samples of fertile and subfertile men, as well as epididymal sperm from *Prm2* mice. Representative images of PTMs, demonstrating significant differences, display fluorescence markings for cell nuclei in blue (Hoechst) and for the respective PTM in red (upper double row = H3K4me3, middle double row = H4K5ac, and lower double row = H4K12ac), with overlays of these two markings appearing pink. Sperm heads marked with white arrows (H3K4me3) exhibited red staining but lacked blue nuclear staining. Identification of these structures as sperm heads was confirmed through the overlay of fluorescence marking (FM) and light microscopic images (LM), revealing the associated flagella. The analysis involved five animals per genotype for *Prm2* mice (n = 5), with a minimum of 50 sperm per animal, and human sperm from five individuals, also with a minimum of 50 sperm per individual; scale bar = 20 µm.

**Figure 6.**
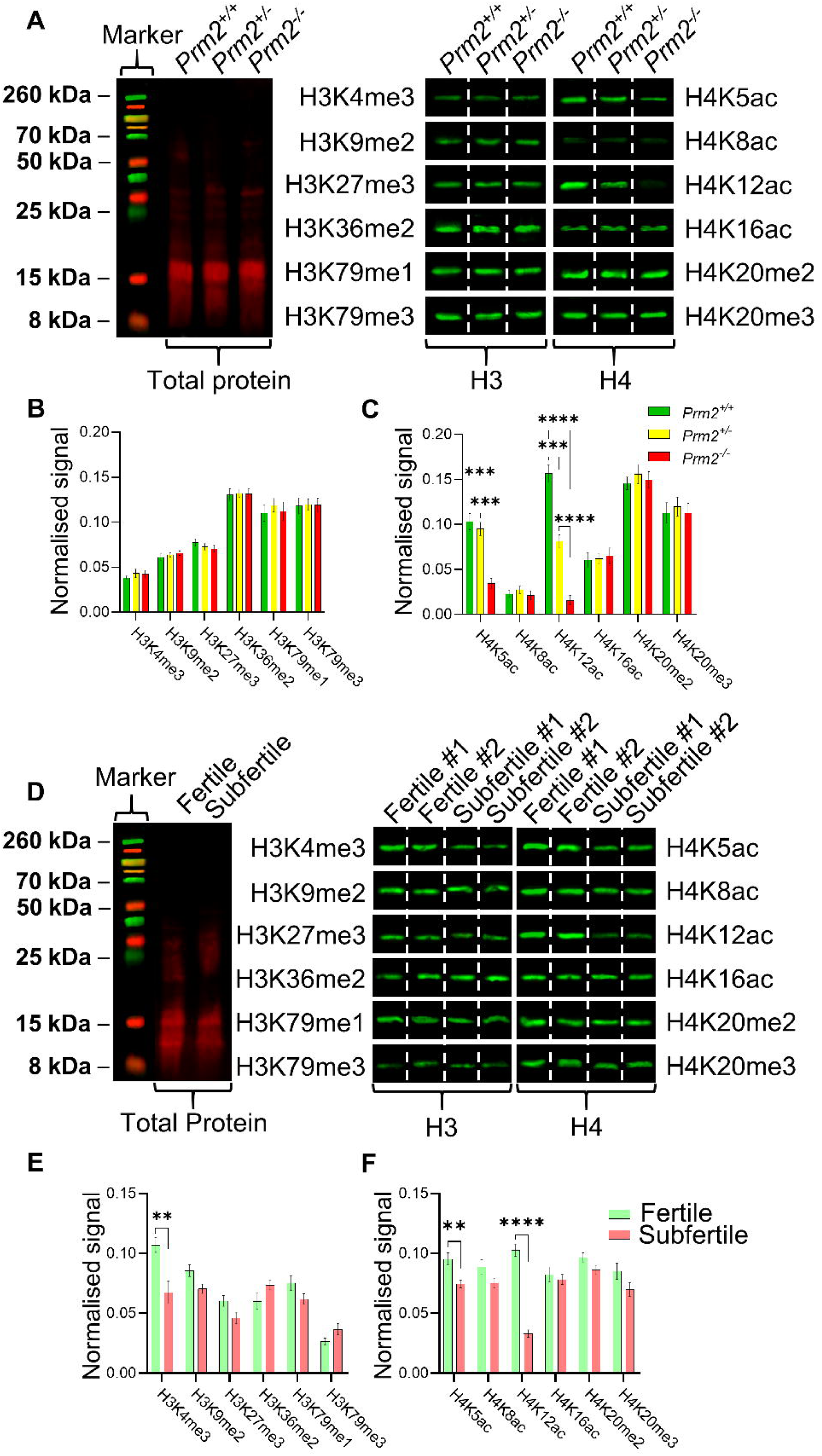
Semi-quantitative comparison of PTM levels for murine and human sperm. Epididymal sperm of *Prm2* mice (**A-C**) and sperm of fertile and subfertile men (**D-F**) were analysed semi-quantitatively for the selected PTMs of H3 and H4 by immunoblotting. **A** and **D**: The representative band pattern of the total protein (left) and the band specific for a PTM (right) of the different genotypes or for human sperm samples are shown, whereby the ejaculate samples used were divided into two fertile and two subfertile sperm samples. Based on the signal values determined for the total protein, signal values of the PTMs were normalised and compared between the genotypes/groups. Mean values of the normalised signal values with s.e.m. for H3 (**B** and **E**) and H4 (**C** and **F**) are shown. Statistical analysis and determination of significant differences were carried out using multiple t-tests, whereby the p-values for multiple testing were adjusted using the Holm-Sidak method. ***Prm2* mice**: <u>Histone H3</u>: no significant differences; <u>Histone H4</u>: H4K5ac *Prm2^+/+^*/ *Prm2^-/-^* *** p = 0.000191; H4K5ac *Prm2^+/-^*/ *Prm2^-/-^* *** p = 0.000172; H4K12ac *Prm2^+/+^*/ *Prm2^-/-^* **** p = <0.000001; H4K12ac *Prm2^+/+^*/ *Prm2^+/-^* *** p = 0.000177; H4K12ac *Prm2^+/-^*/ *Prm2^-/-^* **** p = 0.000067; per genotype n = 5. **Men:** <u>Histone H3</u>: H3K4me3 ** p = 0.0024; <u>Histone H4</u>: H4K5ac ** p = 0.0028; H4K12ac **** p < 0.000001; per group n = 10.

## Discussion

Post-translational modifications of histones are considered to be an important epigenetic mechanism that regulates various cellular processes during spermatogenesis and in mature spermatozoa (Govin *et al*., 2004; Rathke *et al*., 2014; Luense *et al*., 2016). Several informative studies have investigated specific histone PTMs in male germ cells, using antibody-based approaches, while more recent research has utilized mass spectrometry to identify histone PTMs in mature sperm from mice or men (Brunner *et al*., 2013; Luense *et al*., 2016; Schon *et al*., 2019). However, to date, no study has compared mouse and human sperm with aberrant protamine ratio and impaired sperm protamination. This study conducted such a comparison, offering deeper insights into the complex relationship among *Prm2* levels, histone expression patterns, and post-translational modifications (PTMs) of core histones H3 and H4 in murine and human sperm. The fact that impaired sperm protamination or aberrant protamine ratio is associated with male subfertility is well known from numerous studies in the past (Oliva, 2006; Boissonnas *et al*., 2013; Francis *et al*., 2014; Ni *et al*., 2016). Despite years of research, a large part of the molecular processes behind this disorder remain unknown. However, previous studies using a *Prm2*-deficient mouse line, which is also used in this study, have provided insights into defective protamination, demonstrating its suitability as a model for deregulated protamine expression observed in subfertile men (Schneider *et al*., 2016; Schneider *et al*., 2020; Arévalo *et al*., 2022).

The decrease of *Prm2* in heterozygous mice was also found in human subfertile sperm. The reduced relative expression of *PRM2* with unchanged *PRM1* expression in sperm of subfertile men not only confirms the assumption of a link between aberrant protamine ratio and fertility disorders, but is also consistent with the protamine ratios of *Prm2^+/-^* mice. Since the transcripts of protamines are already present in round spermatids of the mid-stages of spermatogenesis, it can be assumed that decreased *Prm2* levels or *Prm2*-deficiency have an influence from this step onwards during spermatogenesis, directly before the start of the histone-protamine exchange (Steger *et al*., 2000; Steger *et al*., 2001). For *Prm2* mice, previous studies did not reveal any noticeable effects on testicular sperm, as they were phenotypically identical across genotypes (Schneider *et al*., 2016; Schneider *et al*., 2020). Furthermore, no differences were found regarding the presence of the core histones H3 and H4 in testis, as both the H3 levels and corresponding immunohistochemical staining for H3 and H4 were comparable across genotypes (Schneider *et al*., 2016). Similarly, immunostaining of *Prm2* mice testis for PTMs of histones H3 and H4 showed no difference between genotypes (*Wt, Prm2^+/-^* and *Prm2^-/-^*) and were consistent with the results of previous studies evaluating histone PTMs in testis of ICR and C57BL/6J mice (Song *et al*., 2011; Tatehana *et al*., 2020). Thus, there seems to be no impact of *Prm2* loss on histones and their PTMs in murine testis. With regard to histone abundance, similar observations were also reported for the *Prm1* mouse model, in which no differences could be detected in testis (Merges *et al*., 2022). The lack of histone (PTMs) staining after step 12 of spermiogenesis, as observed in this study, confirms the successful histone-to-protamine exchange. Furthermore, the absence of staining against H3 and H4 in testicular sperm could be attributed to either inadequate decondensation or insufficient antigen unmasking in fixed spermatids, leading to the inaccessibility of antibodies to their target epitopes. Another possibility is the displacement of H3 and H4 during spermiogenesis to levels below the detection threshold (Ueda *et al*., 2017; Yoshida *et al*., 2018; Rathje *et al*., 2019). Nevertheless, these findings point out the limitations of the staining method, since more sensitive techniques like mass spectrometry have revealed that testicular sperm contain histones with PTMs, although in minimal quantities (Luense *et al*., 2016; Torres-Flores and Hernández-Hernández, 2020).

It has recently been demonstrated that, contrary to initial assumptions, histone-to-protamine exchange is not entirely finalized upon sperm exit from the testis, and histone replacement continues during sperm transit through the epididymis (Yoshida *et al*., 2018). Previous studies showed, that infertility in *Prm2*-deficient mice can be linked to defects in epididymal sperm morphology, motility, and DNA integrity, suggesting the possibility of histone-related effects in epididymal sperm for this study as well (Schneider *et al*., 2016; Schneider *et al*., 2020). A determination of the abundance of H3 and H4 in epididymal sperm of *Prm2* mice showed different levels when comparing genotypes, so that an indirect influence of the different *Prm2* levels on the abundance of core histones can be assumed. The increased abundance of H3 found in heterozygous males could be understood as compensation for the lower amount of protamines (Schneider *et al*., 2016). Whereas the higher level of H4 in sperm of *Prm2*-deficient animals compared to *Wt* and *Prm2^+/-^* is more likely due to the massively disturbed histone-protamine exchange. This incomplete exchange with an increased retention of H4 results in less condensed and more fragile chromatin, making sperm DNA more prone to fragmentation (González-Marín *et al*., 2012). As increased histone abundance is known to be connected to DNA fragmentation in sperm, it was not surprising to find nearly completely degraded DNA in *Prm2*-deficient sperm (Simon *et al*., 2011; Schneider *et al*., 2016; Pandya *et al*., 2024). Fragmentation of DNA in *Prm2*-deficient mice is attributed to the consequences of oxidative stress and in detail to a strong decline in reactive oxygen species (ROS) scavenger proteins SOD1 (superoxide dismutase 1) and PRDX5 (peroxiredoxin 5) and a corresponding increase in ROS levels (Schneider *et al*., 2020). Given that sperm from infertile men are also characterised by DNA fragmentation, some studies have also found higher levels of histone retention in human sperm (Aitken and de Iuliis, 2010; Mohanty *et al*., 2016), although this could not be confirmed in our study. Nevertheless, it can be assumed that the investigated group of subfertile human spermatozoa has a higher proportion of DNA fragmentation than the fertile group, as the reduced *Prm2* level impedes sufficient hypercondensation and protection of the paternal genome. In addition, the undisputed detection and abundance of H3 and H4 in human sperm is consistent with the results of other studies and supports the comparability of murine and human histone retention in sperm (Yamaguchi *et al*., 2018; Yoshida *et al*., 2018).

As numerous studies have shown, the PTM profile of retained H3 and H4 is comparable across fertile murine and human sperm. Thus, it remains of interest to investigate these profiles when there are changes in sperm quality (Brykczynska *et al*., 2010; Hammoud *et al*., 2011; Erkek *et al*., 2013; Krejčí *et al*., 2015; Luense *et al*., 2016; Schon *et al*., 2019). In our study we were able to confirm the similarities between the methylation and acetylation of the lysine residues of H3 and H4 by means of a comprehensive mass spectrometric analysis. For *Prm2* mice, there were no significant differences for histone H3 except for the significantly higher levels of H3K9me3 in *Prm2^+/-^*. This increase could also be the result of an attempt to compensate for reduced *Prm2* level, as H3K9me3 is associated with heterochromatin and contributes to genome stability (Peters *et al*., 2001; Sullivan and Karpen, 2004). In contrast, the methylation of H3K9 in sperm of fertile and subfertile men was unchanged. Interestingly, a significant decrease in the histone variant H3.3 (H3.3K27K36K37unmod) was found in the mass spectrometric analysis of H3 for subfertile sperm. These reduced H3.3 levels already showed an effect on sperm quality in corresponding knockout models and are presumably also connected to reduced human sperm quality (Yuen *et al*., 2014; Tang *et al*., 2015). The increased levels of euchromatin-associated PTM H3K23ac and heterochromatin marker H3K27me2 in subfertile sperm indicate disrupted epigenetic regulation of germ cell differentiation during spermatogenesis, whereby the exact mechanisms need to be clarified in future studies (Sendler *et al*., 2013; Chioccarelli *et al*., 2020; Grant *et al*., 2022; Kumar *et al*., 2024). Unfortunately, due to low abundance and short retention time of H3K4me3 in mammalian cells, our mass spectrometric analysis could not reliably detect the prominent PTM in murine or human sperm (Garcia *et al*., 2007; Lesch and Page, 2014). Nevertheless, H3K4me3, as one of the best analysed PTMs in general, was part of the follow-up experiments using Western blot and immunofluorescent staining, where reduced levels of H3K4me3 could be found in sperm of subfertile men. This decrease could have an influence on the epigenetic regulation of important developmental genes, as H3K4me3 is associated with CpG-rich promoters of such genes (Brykczynska *et al*., 2010; Yamaguchi *et al*., 2018). Since epididymal sperm from *Prm2* mice did not show any differences for H3K4me3 comparing the genotypes, a non-protamine-associated mechanism for the redistribution of H3K4me3 levels or the other affected PTMs of H3 (H3K27me2 and H3K23ac) can be assumed. The additional analysis of H3K9me2, H3K27me3, H3K36me2, H3K79me1 and H3K79me3 did not reveal significant differences in *Prm2* mice or human samples, suggesting these modifications are not impacted by the consequences of reduced *Prm2* levels.

For H4 PTMs, our mass spectrometric results showed that acetylations of peptide H4_4-17 (H4ac) exhibited comparable intraspecific differences for murine and human samples. The significantly lower levels of H4ac in *Prm2^+/-^* compared to *Wt* are extremely similar to those of subfertile human spermatozoa and therefore suggest a possible misregulation of H4ac as a result of altered *Prm2* levels in both species. Together with the antibody-based results, these reduced H4ac levels could be attributed to H4K5ac and H4K12ac, both modifications that are involved in the process of chromatin remodelling during spermiogenesis (Kim *et al*., 2014; Chen *et al*., 2021; de la Iglesia *et al*., 2022). The assumption that reduced H4ac levels in sperm are associated with reduced chromatin condensation and, consequently, abnormal spermiogenesis and sub-to infertility is consistent with the findings of other studies on bull and human sperm (Paradowska *et al*., 2012; Schon *et al*., 2019; Ugur *et al*., 2019). In combination with the *Prm2* mouse model, the comparable results found in this study suggest that reduced levels of *PRM2* in human sperm are also linked to decreased levels of H4K5ac and H4K12ac. This implies that a defective protamine ratio may have far-reaching epigenetic consequences. That such decreased H4ac level are also found in morphologically normal sperm, such as *Prm2^+/-^* sperm, was confirmed by our results and is supported by other studies with human sperm (Schon *et al*., 2019). The observed decrease in acetylation of H4K5ac and H4K12ac in both mice and men could be a consequence of the aforementioned secondary effects such as compromised DNA integrity or also a consequence of misregulated histone acetyltransferases (HATs) or histone deacetylases (HDACs). Especially in the sperm of *Prm2^-/-^*, the previous findings suggest that the almost complete loss of H4ac is a result of a self-destruction cascade of the sperm initiated by faulty DNA protamination, as described in previous studies (Schneider *et al*., 2020).

Our findings align with recent studies highlighting the dynamic nature of epigenetic regulation in spermatogenesis and its implications for male fertility (Luense *et al*., 2016; Gold *et al*., 2018; Chioccarelli *et al*., 2020; Tahmasbpour Marzouni *et al*., 2022). In conclusion, our investigations provide valuable insights into the complex interplay among *Prm2* levels, histone levels, and post-translational modifications (PTMs) in the context of male fertility. Reduced *Prm2* levels could be linked to decreased H4ac levels in both mouse and man and suggest a disturbed histone-protamine exchange also on the epigenetic level. Since *Prm2* probably has only an indirect influence on histone PTMs, further studies are needed to uncover the underlying mechanisms and to understand such epigenetic changes in detail.

## Supplementary Data

Supplementary data are available in supplementary information files.

## Data availability

The data underlying this article are available in the article and in its online supplementary material.

## Supporting information

Human mass spec data

Mouse mass spec data H3

Mouse mass spec data H4

## Acknowledgements

This study was supported by a grant from the German Research foundation (DFG) to HS (SCH 503/15-2) and KS (STE 892/14-2). We would like to thank the colleagues of the Clinic and Polyclinic for Urology, Paediatric Urology and Andrology of the University Clinic Giessen and Marburg, in cooperation with Prof. Dr. Adrian Pilatz, Prof. Dr. Hans-Christian Schuppe, under the direction of Prof. Dr Florian Wagenlehner for providing and clinically assessing human samples for this project. In this context, it is noteworthy to mention and thank the Hormone and Fertility Centre of the Ludwig-Maximilians-University Munich, where further human sample material was provided through collaboration with Prof. Dr Nina Rogenhofer. In addition, we thank Barbara Fröhlich, Kerstin Wilhelm, Tania Bloch, Gaby Beine, Angela Egert, Andrea Jäger, for excellent technical assistance.

## Authors’ roles

Conceptualization: AK, KS, HS.; Methodology: AK, IF, AI; Formal analysis: AK, SS, GEM, ACF, IF, AI, HS, KS; Investigation: AK, IF; Resources: KS, HS; Data curation: AK; Writing - original draft: AK; Writing - review & editing: AK, ACF, KS, HS; Visualization: AK; Supervision: KS, HS; Project administration: KS; Funding acquisition: KS, HS.

