## Supplemental figures for "An aberrant protamine ratio is associated with decreased H4ac levels in murine and human sperm"

### Supplementary Data

**Supplementary figure S1: Immunofluorescent detection of the nuclear histones H3 and H4 in murine and human sperm.**

Fluorescence microscopic, representative images showed cell nuclei by Hoechst (blue) and histones H3 (**A** and **C**) and H4 (**B** and **D**) in red for murine (**A** and **B**) and human (**C** and **D**) sperm. When nuclear staining and antibody staining overlapped, these areas showed up as pink. Overlay of fluorescent labelling (FM) and light microscopic image (LM) also revealed the associated flagella of sperm heads. For evaluation of the staining, five animals (n = 5) and at least 50 spermatozoa per animal were counted per genotype; scale bar = 20 µm.

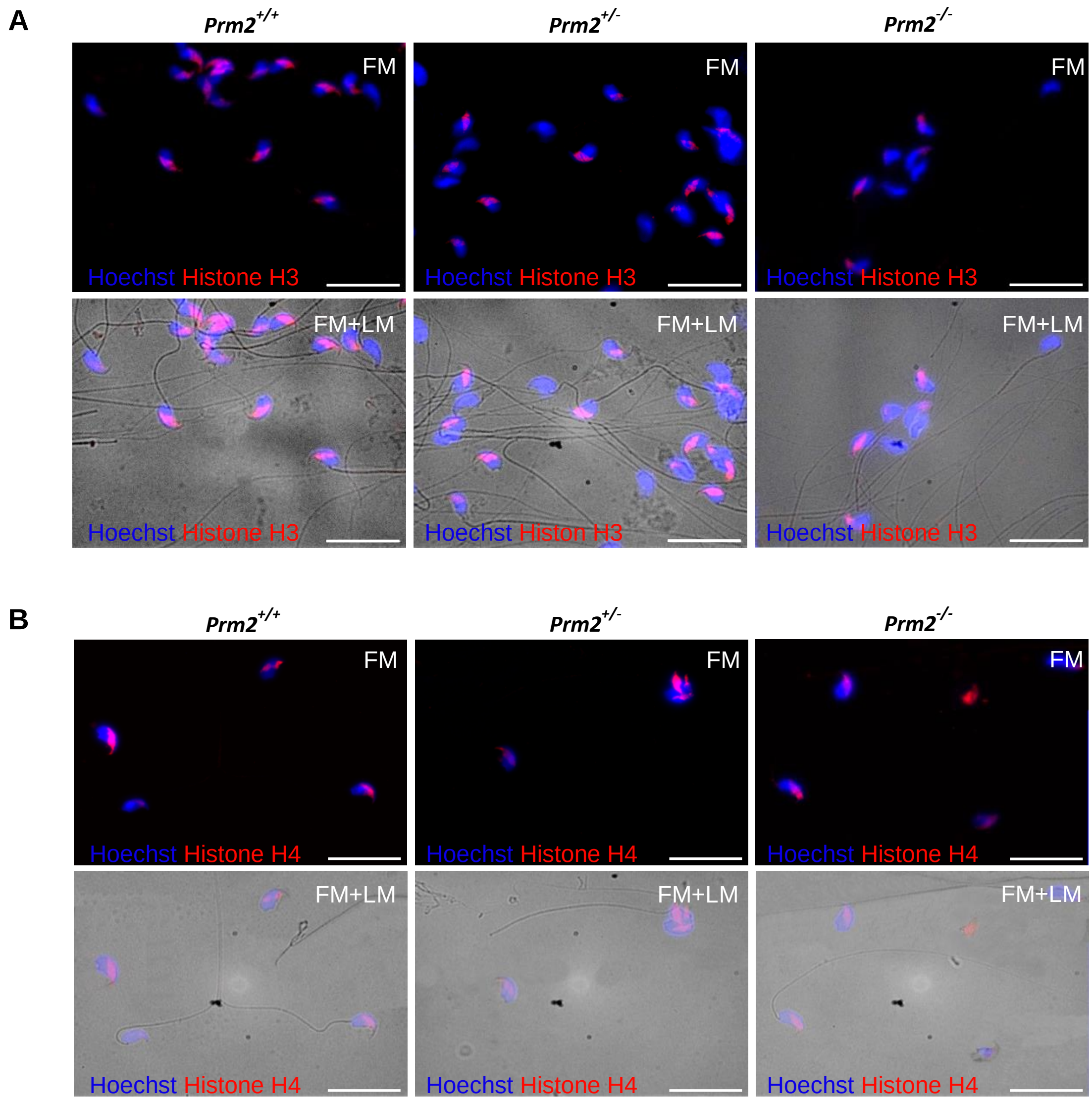

C

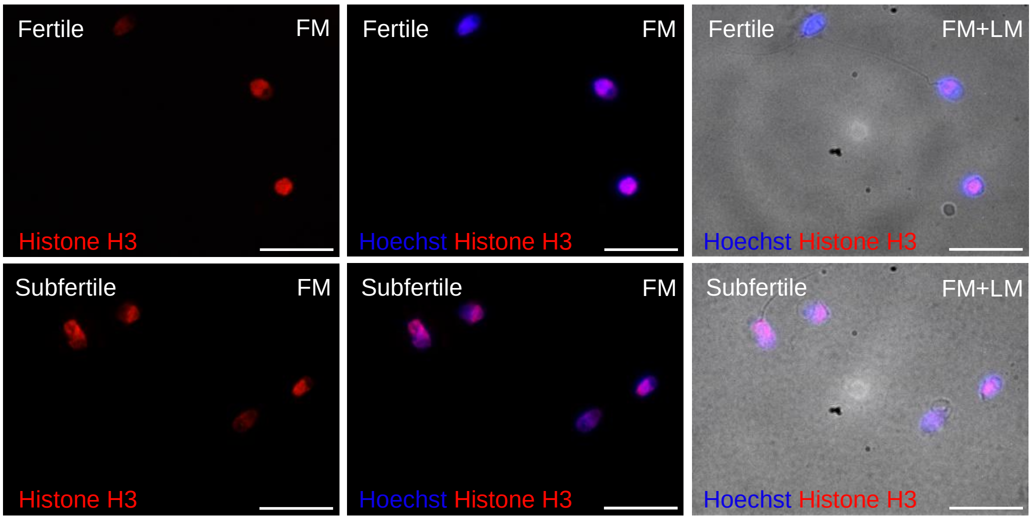


D

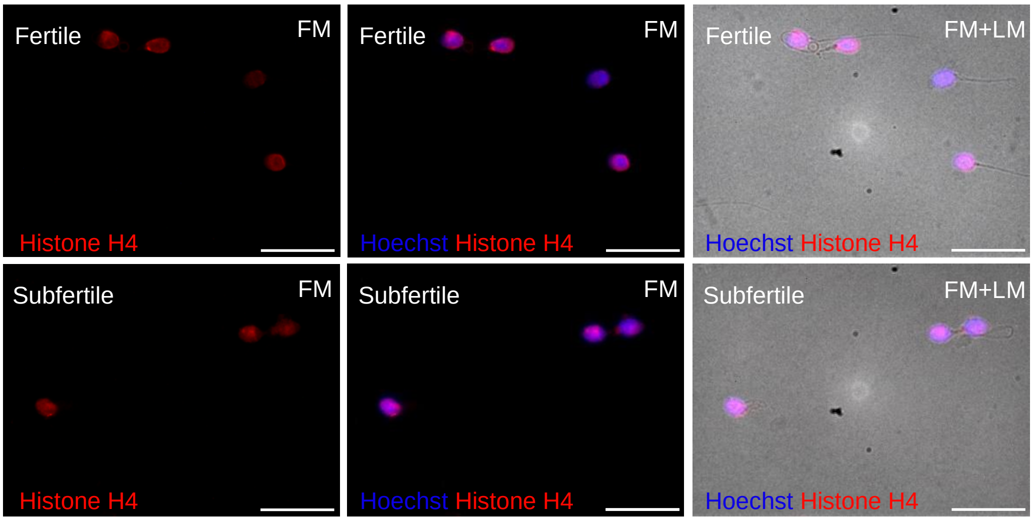


**Supplementary Figure S2: Histone H3 PTM expression pattern in the testis of *Prm2* mice.**

Shown are representative images of the immunohistochemical detection of the selected PTMs of histone H3 in the testis of *Prm2* mice, as well as a schematic representation of the stage-specific expression pattern. A half overlay of cell types in the schematic indicates weak or partial staining (figure modified from (Russell *et al.*, 1993; Endo *et al.*, 2015)). Positive cells showed red staining and the remaining nuclei were blue by counterstaining with haematoxylin. The stages of spermatogenesis are labelled with Roman numerals. **A-D**: H3K4me3, **E-H**: H3K9me2, **I-L**: H3K27me3, **M-P**: H3K36me2, **Q-T**: H3K79me1 and **U-X**: H3K79me3. Cell types: Spermatogonia type A (A); leptotene spermatocytes (L), zygotene spermatocytes (Z), pachytene spermatocytes (P), diplotene spermatocytes (D), secondary spermatocytes in meiosis (m2Om/M), 1-8 round spermatids (RS), 9-16 elongating (ES)/ elongated (ET) spermatids. Staining of five animals per genotype was analysed (n = 5); scale bar = 50 µm.

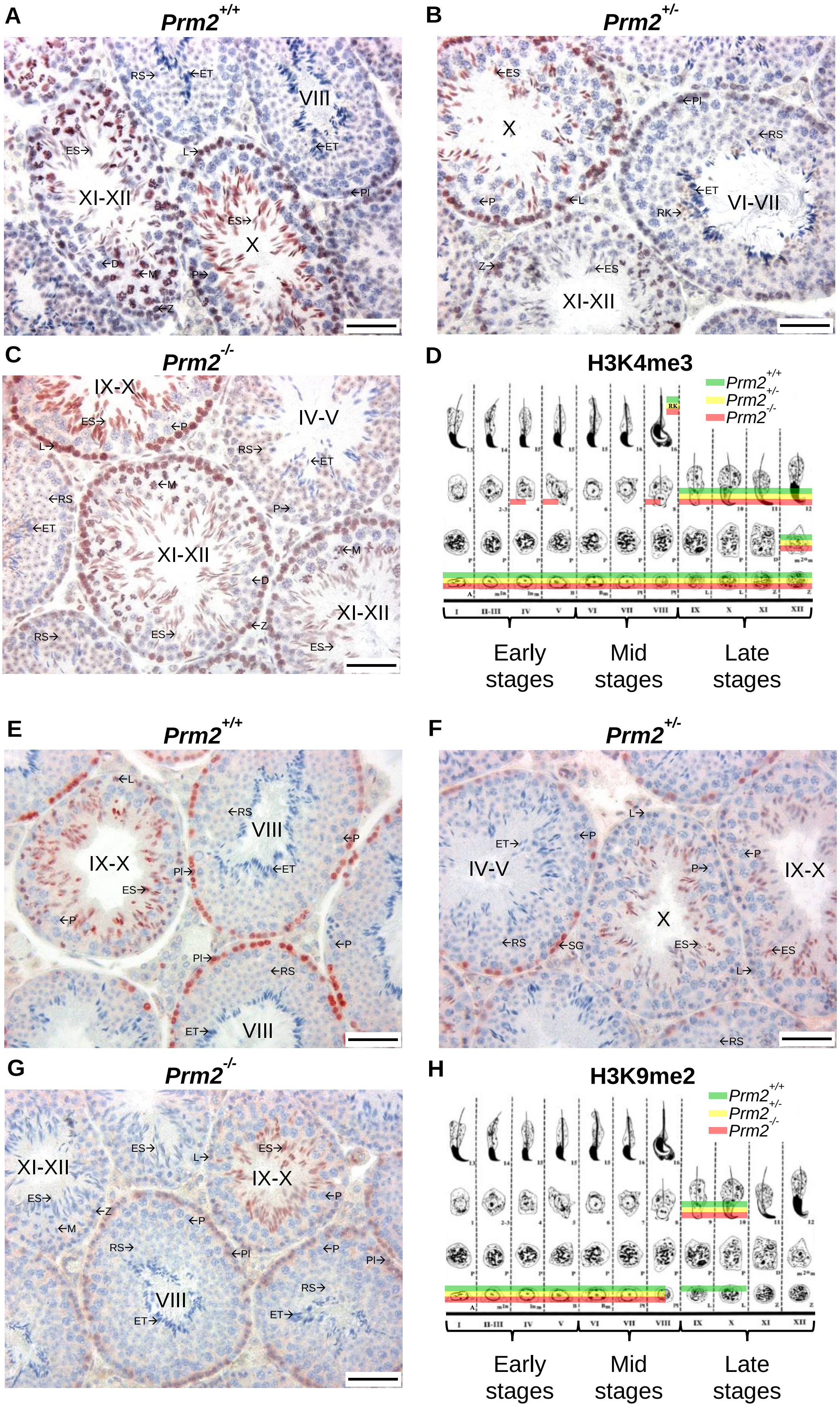

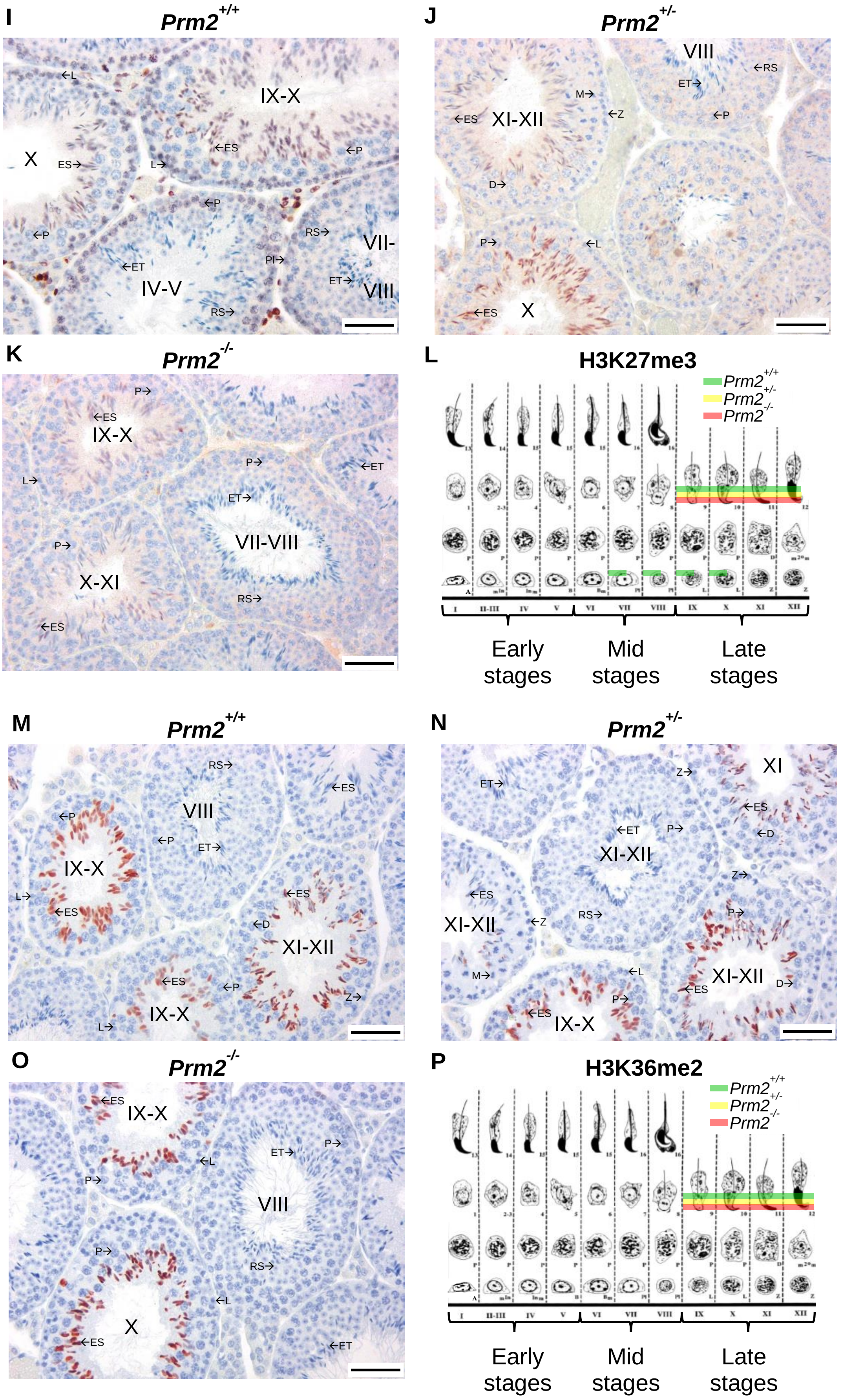

I
Prm2+/+
IX-X
X
VII-
VIII
IV-V
J
Prm2+/-
VIII
XI-XII
X
K
Prm2-/-
IX-X
VII-VIII
X-XI
L
H3K27me3
Prm2+/+
Prm2+/-
Prm2-/-
Early stages
Mid stages
Late stages
M
Prm2+/+
VIII
IX-X
XI-XII
IX-X
N
Prm2+/-
XI
XI-XII
XI-XII
XI-XII
IX-X
O
Prm2-/-
IX-X
VIII
X
P
H3K36me2
Prm2+/+
Prm2+/-
Prm2-/-
Early stages
Mid stages
Late stages

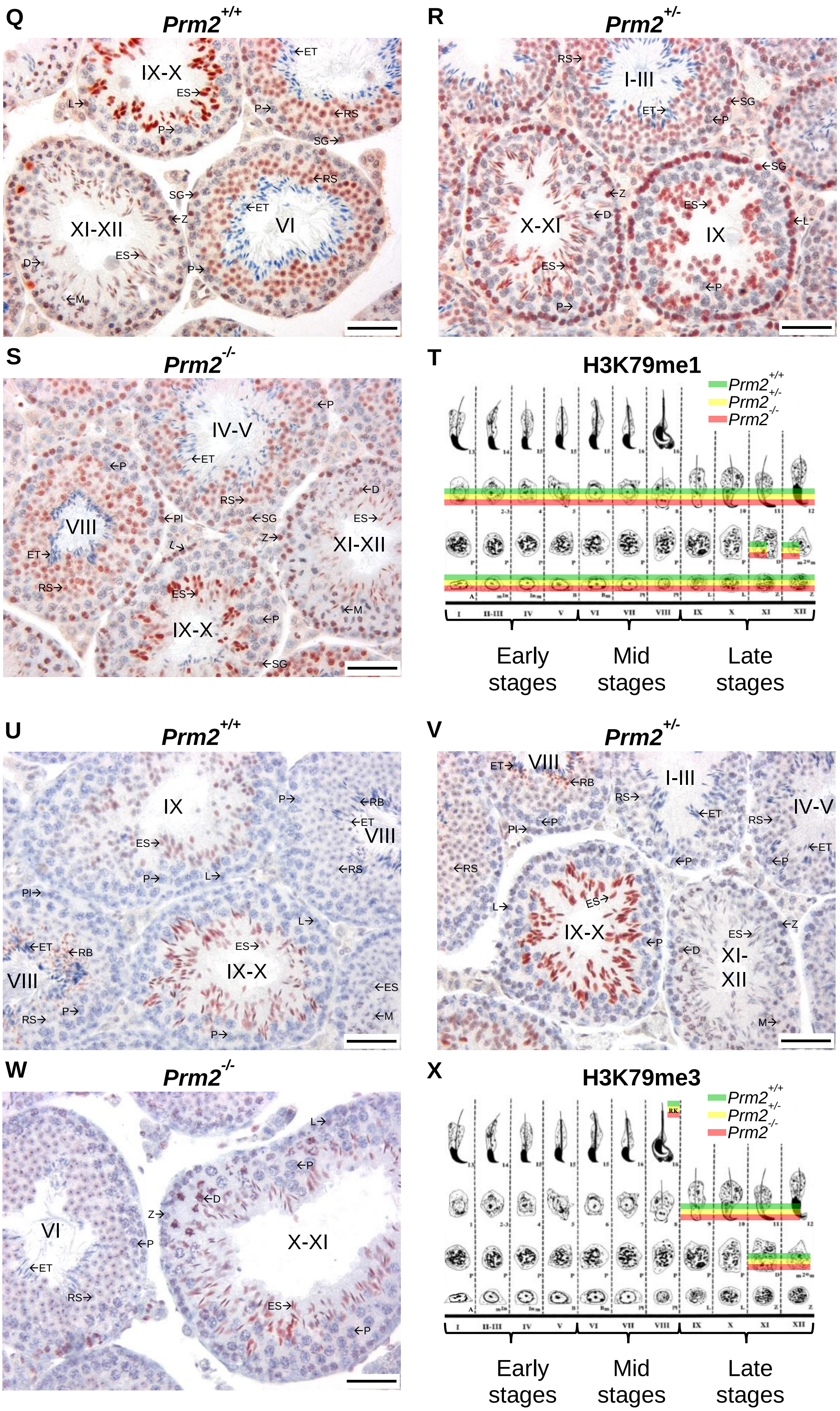

Q
Prm2+/+
IX-X
XI-XII
VI
R
Prm2+/-
I-III
X-XI
IX
S
Prm2-/-
IV-V
VIII
XI-XII
IX-X
T
H3K79me1
Prm2+/+
Prm2+/-
Prm2-/-
I
II-III
IV
V
VI
VII
VIII
IX
X
XI
XII
Early stages
Mid stages
Late stages
U
Prm2+/+
IX
VIII
IX-X
V
Prm2+/-
VIII
I-III
IV-V
IX-X
XI-XII
W
Prm2-/-
VI
X-XI
X
H3K79me3
Prm2+/+
Prm2+/-
Prm2-/-
I
II-III
IV
V
VI
VII
VIII
IX
X
XI
XII
Early stages
Mid stages
Late stages

**Supplementary Figure S3: Expression of selected PTMs of H4 in the testis of *Prm2* mice.**

Representative images depicting the immunohistochemical detection of specific PTMs of histone H4 within *Prm2* mouse testicular tissue and a schematic diagram, adapted from (Russell et al. 1993; Endo et al. 2015), that illustrates stage-specific patterns of expression are presented. The schematic incorporates semi-overlays denoting weak or partial staining. Positively stained cells are visualized in red, while the remaining nuclei are counterstained in blue with haematoxylin. Roman numerals are utilized to label the stages of spermatogenesis. **A-D**: H3K4me3, **E-H**: H3K9me2, **I-L**: H3K27me3, **M-P**: H3K36me2, **Q-T**: H3K79me1 and **U-X**: H3K79me3. Cell types: Spermatogonia type A (A); leptotene spermatocytes (L), zygotene spermatocytes (Z), pachytene spermatocytes (P), diplotene spermatocytes (D), secondary spermatocytes in meiosis (m2Om/M), 1-8 round spermatids (RS), 9-16 elongating (ES)/ elongated (ET) spermatids. Staining of five animals per genotype was analysed (n = 5); scale bar = 50 µm.

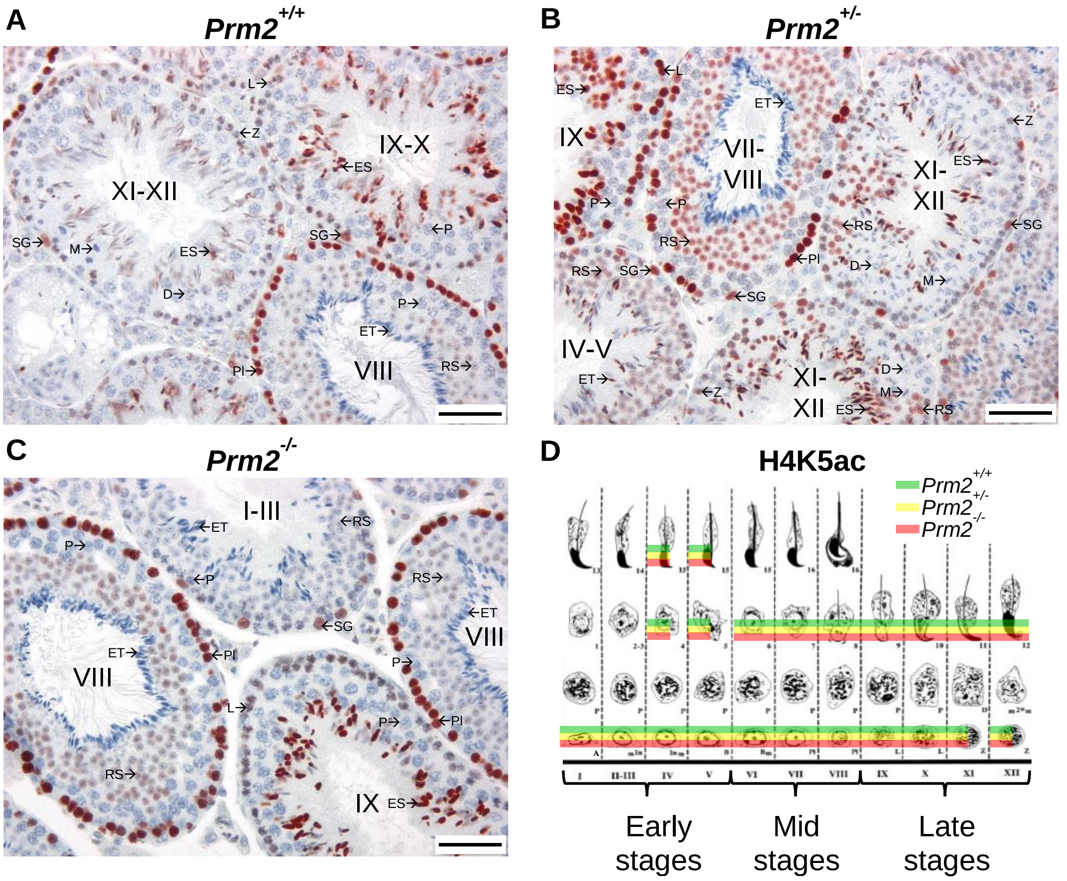

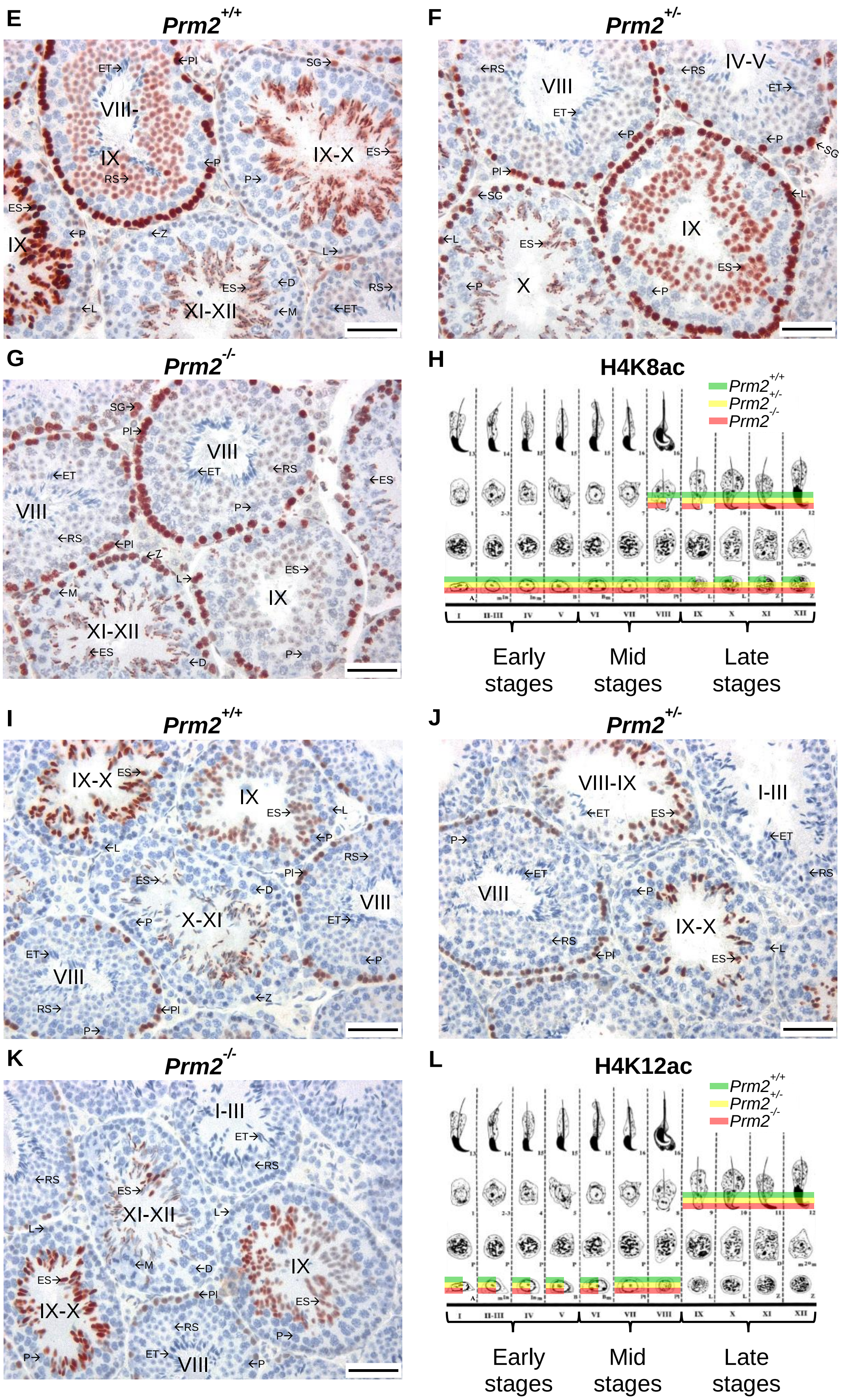
E
Prm2+/+
ET→
←PI
SG→
VIII-
IX
RS→
←P
IX-X
ES→
P→
ES→
IX
←P
←Z
L→
ES→
←D
RS→
←L
XI-XII
←M
←ET
F
Prm2+/-
←RS
VIII
←RS
IV-V
ET→
ET→
←P
←P
←SG
PI→
←SG
←L
←L
ES→
IX
ES→
←P
X
←P
G
Prm2-/-
SG→
PI→
VIII
←ET
←RS
←ET
←ES
P→
VIII
←RS
←PI
←Z
L→
ES→
←M
IX
XI-XII
←ES
←D
P→
H
H4K8ac
Prm2+/+
Prm2+/-
Prm2-/-
Early stages
Mid stages
Late stages
I
Prm2+/+
IX-X
ES→
IX
ES→
←L
←L
←P
RS→
ES→
PI→
←D
VIII
←P
X-XI
ET→
ET→
←P
VIII
RS→
←PI
←Z
P→
J
Prm2+/-
VIII-IX
I-III
←ET
ES→
←ET
P→
←ET
←RS
←P
VIII
IX-X
←RS
←PI
←L
ES→
K
Prm2-/-
I-III
ET→
←RS
←RS
ES→
L→
XI-XII
L→
←M
←D
IX
ES→
←PI
IX-X
ES→
←RS
P→
ET→
VIII
←P
P→
L
H4K12ac
Prm2+/+
Prm2+/-
Prm2-/-
Early stages
Mid stages
Late stages

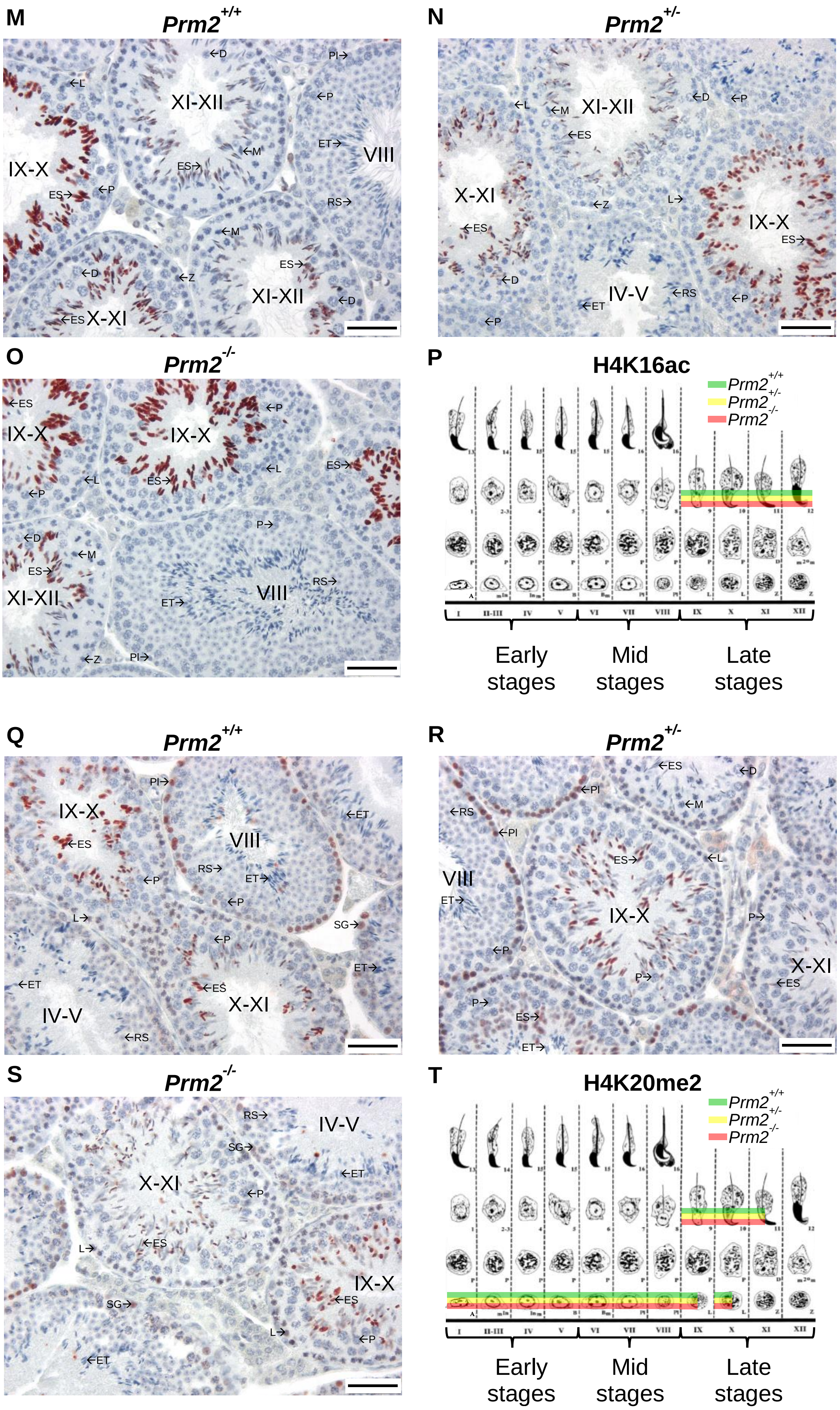
M
Prm2+/+
N
Prm2+/-
O
Prm2-/-
P
H4K16ac
Prm2+/+
Prm2+/-
Prm2-/-
Early stages
Mid stages
Late stages
Q
Prm2+/+
R
Prm2+/-
S
Prm2-/-
T
H4K20me2
Prm2+/+
Prm2+/-
Prm2-/-
Early stages
Mid stages
Late stages

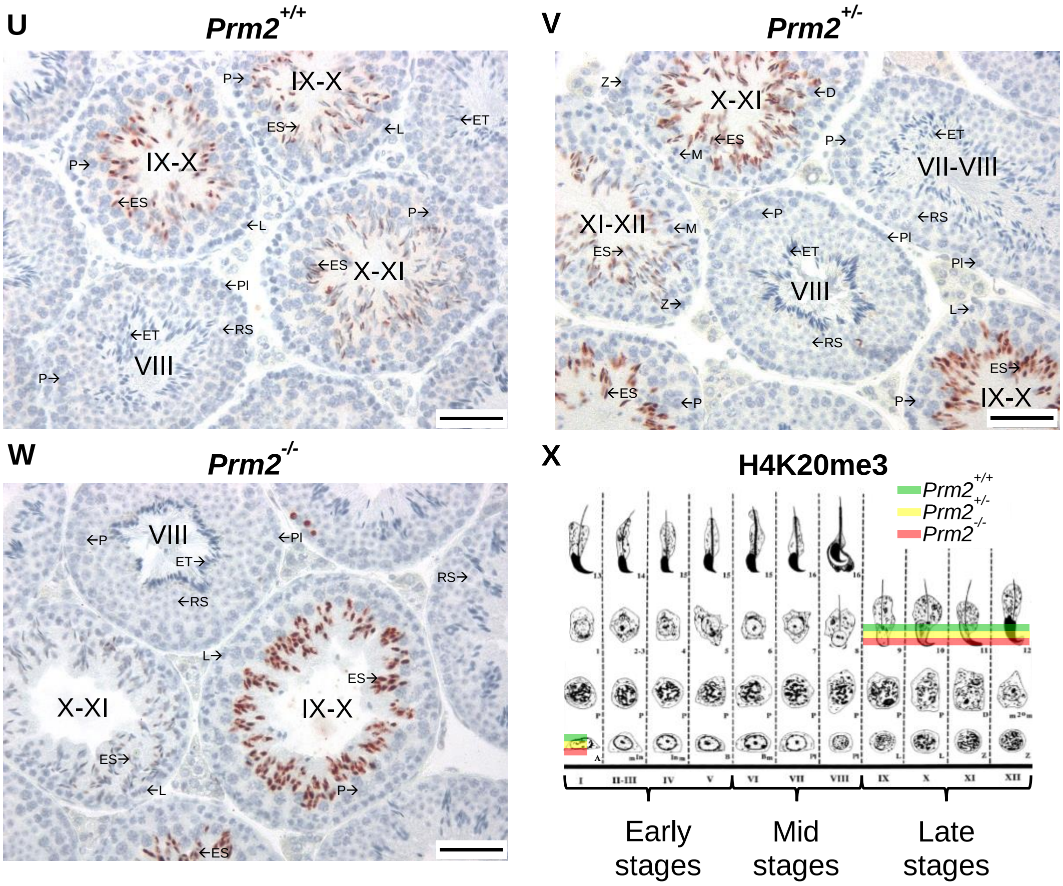


**Supplementary Figure S4: No significant visual differences for a large proportion of PTMs from H3.**

The presented figures illustrate the representative immunofluorescent detection of different PTMs of histone H3 in sperm from both mice and men that showed no significant differences. The fluorescence microscopy images (FM) display blue nuclear staining and red staining of the respective PTM, resulting in a pink appearance upon overlap. By merging FM and light microscopy (LM) images, sperm with typically unmarked flagella and those lacking blue nuclear staining were identified. White arrows point to the sperm heads with intense H3K27me3 that lack nuclear staining. In total, the analysis involved staining 50 sperm from five animals for each genotype and five men per group(n = 5), each with a minimum of 50 sperm analysed per individual; scale bar = 20 µm.

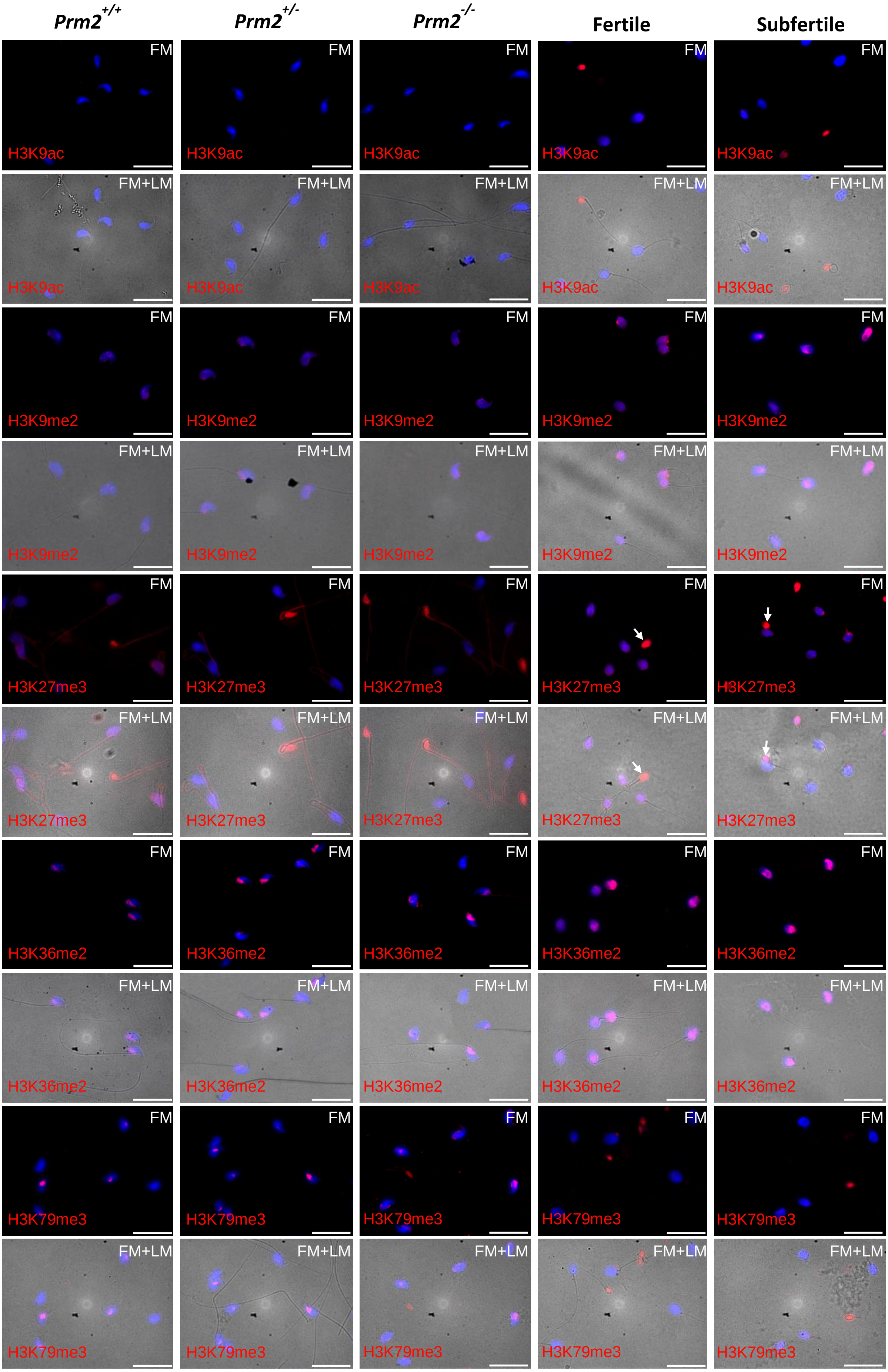

Prm2+/+
Prm2+/-
Prm2-/-
Fertile
Subfertile
FM
FM+LM
H3K9ac
H3K9me2
H3K27me3
H3K36me2
H3K79me3

**Supplementary Figure S5: Absence of significant visual changes for certain acetylations and methylations of H4.**

Immunofluorescent staining against some of the PTMs of H4 on sperm from mice and men was used to assess their visible expression. In the following, only those images are shown in which there were no significant differences between the comparison groups. Fluorescence microscopic images (FM) show a blue nuclear staining and a red staining of the respective PTM, which appears pink when overlapping. Using the overlay of FM and light microscopy (LM) images, sperm with typically unlabelled flagella and those without blue nuclear staining could be identified. White arrows indicate sperm heads with bipartite staining for H4K16ac. Analysis was performed with at least 50 spermatozoa from five animals for each genotype and five males per group (n = 5), also analysing at least 50 spermatozoa per individual; scale bar = 20 µm.

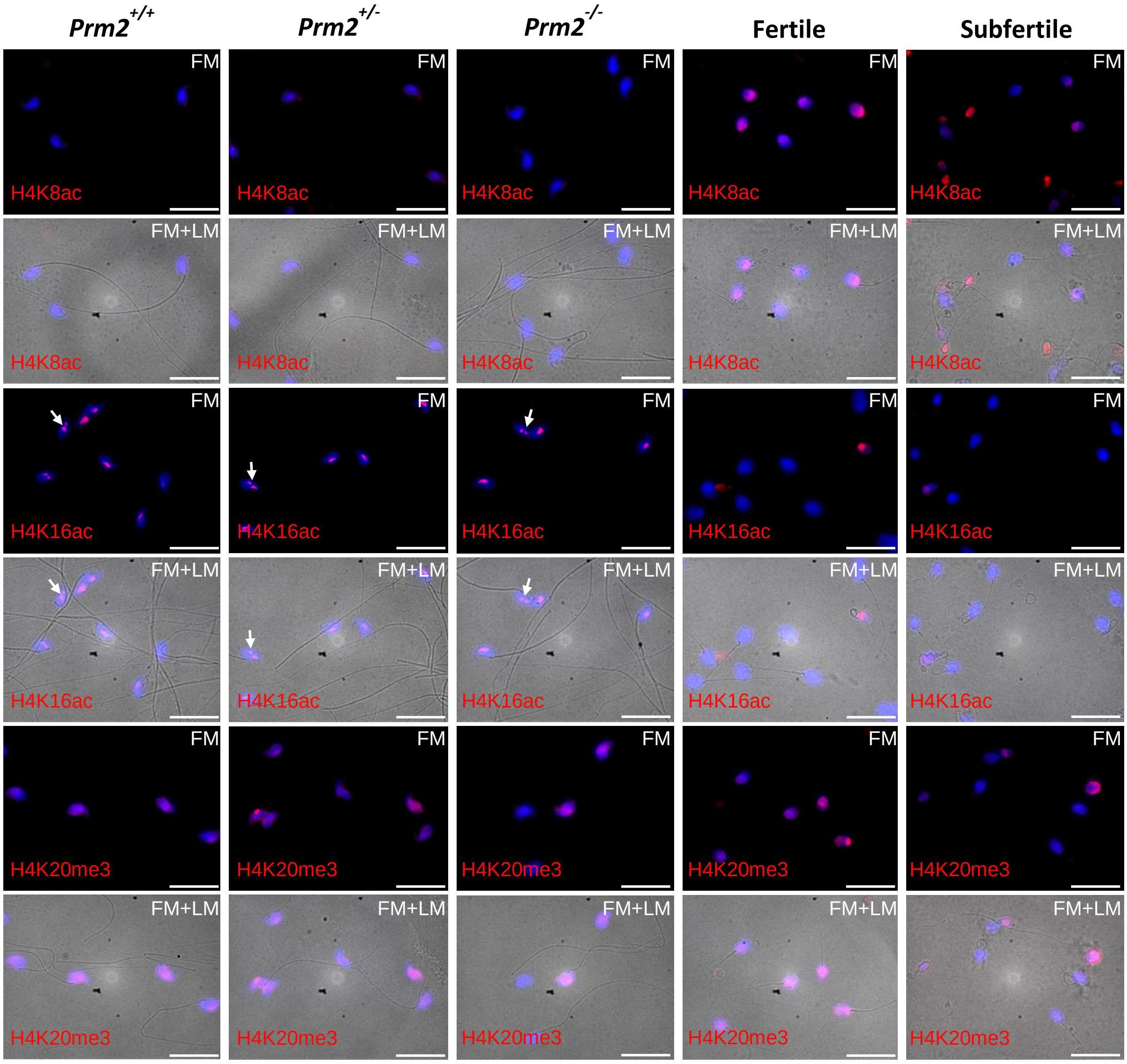

**Supplementary Figure S6: Uncropped western blots for *Prm2* mice and human samples.**

The Western blots shown are the uncropped and unprocessed images, which are related to the results presented in Fig.6. H3 and its PTMs are found at approx. 15 kDa and the bands of H4 and its PTMS at approx. 11 kDa, i.e. directly below. Shown are the separate blots for murine and human samples, with the figure for H3K27me3 and H4K20me2 showing results for both.

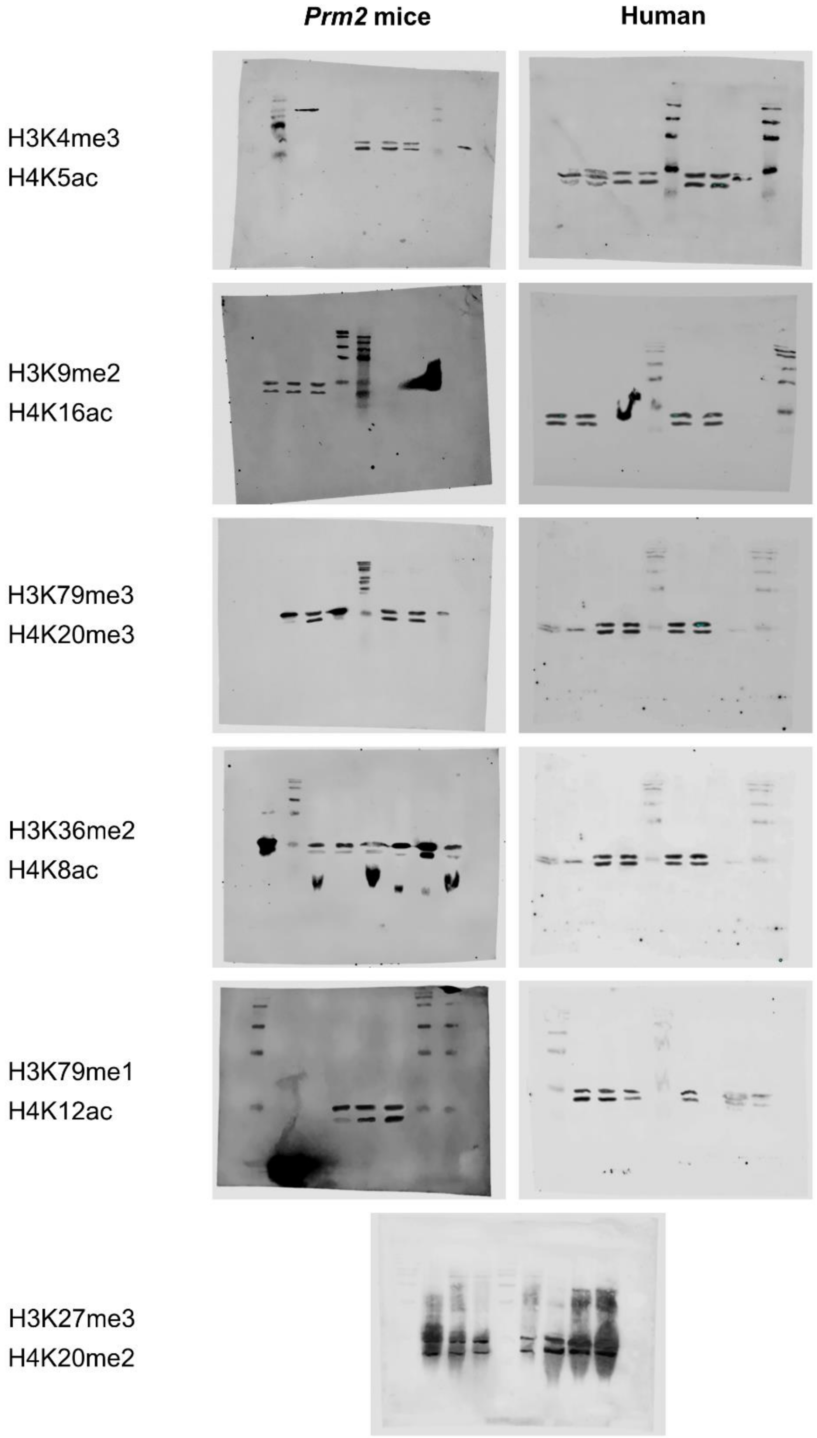

**Supplementary Table SI: Primers used for qPCR.**

| Gene | Accession number | Primers | |
|---|---|---|---|
| | | Forward | Reverse |
| *Prm1* | NM_013637.5 | AGATACCGATGCTGCCGCAGCA | TAGTATTTTTTACACCTTATGGTGTAT |
| *PRM1* | Z46940 | AAGTCGCAGACGAAGGAGG | ATCTCGGTCTGTACCTGGGG |
| *Prm2* (Genotyping) | NM_008933.2 | GAATGAGGAGCCCCAGTGAG (TGCAGCCTCAATCCAGAACC) | TCTGGGCTCAGCCCTTGCCC (TGTAGCCTCTTACGAGAGCAG) |
| *PRM2* | Z46940 | AAGACGCTCCTGCAGGCAC | GCCTTCTGCATGTTCTCTTCCT |
| *Catsper1* | NM_139301.3 | TTGCAGCATTTGCGTGAGTT | GGTGGGGATGATGATTCGGG |
| *ACTB* | NM_001101.5 | GATTCCTATGTGGGCGACGAG | AGGTCTCAAACATGATCTGGGT |

For unprocessed mass spectrometry data see excel files:

Mouse Peptide Quantification_H3.xlsx

Mouse Peptide Quantification_H4.xlsx

Human Peptide Quantification_H3_H4.xlsx
